# Flexible and robust cell type annotation for highly multiplexed tissue images

**DOI:** 10.1101/2024.09.12.612510

**Authors:** Huangqingbo Sun, Shiqiu Yu, Anna Martinez Casals, Anna Bäckström, Yuxin Lu, Cecilia Lindskog, Matthew Ruffalo, Emma Lundberg, Robert F. Murphy

## Abstract

Identifying cell types in highly multiplexed images is essential for understanding tissue spatial organization. Current cell type annotation methods often rely on extensive reference images and manual adjustments. In this work, we present a tool, Robust Image-Based Cell Annotator (RIBCA), that enables accurate, automated, unbiased, and fine-grained cell type annotation for images with a wide range of antibody panels, without requiring additional model training or human intervention. Our tool has successfully annotated over 3 million cells, revealing the spatial organization of various cell types across more than 40 different human tissues. It is open-source and features a modular design, allowing for easy extension to additional cell types.

## Introduction

Cells can be characterized into different types by their molecular, morphological, physiological, and functional properties (1). Different types of cells are precisely localized to allow interactions that give rise to specific environments for physiological functions. Advances in multiplexed tissue imaging now enable the probing of over 50 proteins in a single tissue sample, potentially identifying many different cell types through their proteomic profiles. However, unlike spatial transcriptomics in which thousands of markers are measured, the lower number of markers in spatial proteomics poses challenges for devising generalizable and robust cell annotation frameworks. Traditional approaches rely on manually-specified marker ranges (gates), but they often lack reproducibility and require prior knowledge to choose cell types and ranges(2). Alternatively, supervised machine learning (ML) constructs end-to-end models, but training these ML models requires large reference datasets that are often unavailable and difficult to obtain (3, 4). Unsupervised clustering methods can also be employed for cell labeling, but their performances generally fall short of manual gating or supervised methods.

Furthermore, many methods rely solely on average intensities to annotate cell types, overlooking the spatial information and subcellular protein patterns provided by images. As Rumberger *et al*. demonstrated, the difference in average intensity between positive and negative expressions can be minimal (5). Moreover, current approaches focus more on method development than on creating *ready-to-use* software, leaving users to annotate their own data and retrain models based on published methods.

To address these challenges, we have developed the Robust Image-Based Cell Annotator (RIBCA) for cell type annotation in multiplexed tissue images that is generalizable to new image collections without extra fine-tuning. Instead of a single model, we constructed an ensemble of image-derived models, where each base model is explicitly designed for a subset of protein markers and a subset of cell types of interest (Figure 1**A**). The rationale behind this modular design is to make it compatible with any common antibody panel by matching its markers with one or more base models in this ensemble. Our approach involves cropping whole tissue images into single-cell regions, using imputation models to predict the patterns of any missing markers, predicting with any matched model, and merging the results (Figure 1**B** and **C**). The output is a cell type map, its annotation confidence, and spatial analysis of cell type patterns. Our software has a Napari plugin for interactively validating annotations (Figure S1).

**Fig. 1.**
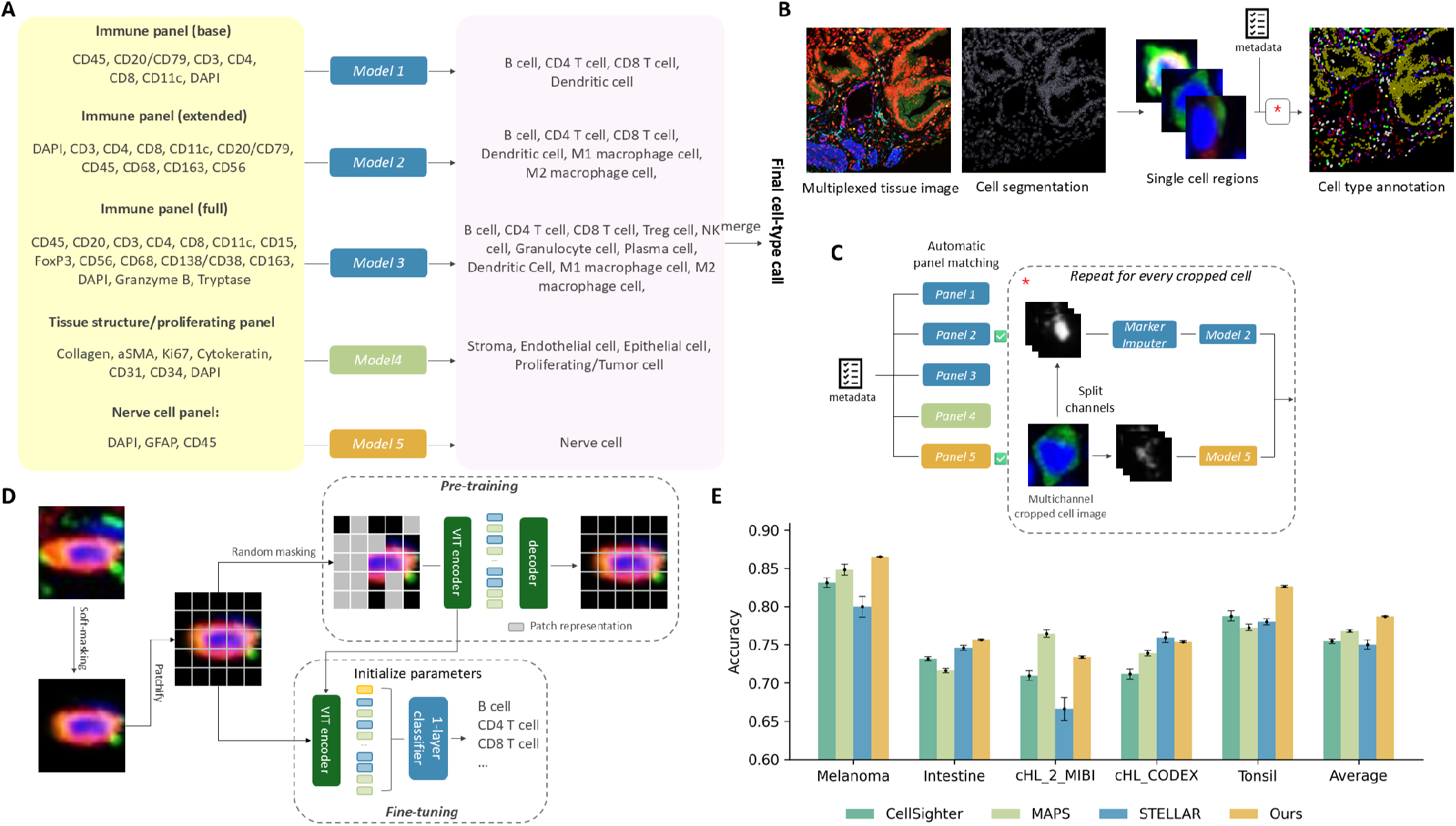
Overview of RIBCA. **A**, Ensemble design for base models to map a subset of image channels to particular cell types. **B**, A multiplexed tissue image and its cell segmentation is cropped into single cell regions to annotate individual cells. **C**, Available channels are checked to determine if they include enough to be acceptable to a given panel (indicated by check marks). In this case, Panels 2 and 5 are selected, Marker Imputers fill in missing markers, cell type models classify the imputed images, and the classification results are merged. **d**, Diagrams of pre-training and fine-tuning Vision Transformer models for cropped cell images. **E**, Average cross-validation accuracies for different methods on five test sets. Error bars indicate ±1 standard deviation from the mean.

## Results

### Overview of RIBCA

Unlike methods that convert pixel intensities into average channel values per cell, image-derived models learn subcellular patterns directly from images and use them to classify. Here we employed the Vision Transformer (ViT) (6), illustrated in Figure 1**D**. We trained individual ViT for five publicly available multiplexed image datasets using 7-fold cross-validation to randomly partition the whole dataset into training and test sets. We then compared our predictions on the test sets with those from three recent methods (see Methods) and calculated the average accuracies (Figure 1**E**). Although no single method outperformed the others for all datasets, ours provides the best overall, and most stable, performance.

Success in training models for each dataset motivated us to develop a generic pre-trained and ready-to-use ensemble model. We began by curating a multiplexed tissue image dataset (referred to as `SPData_1`) containing more than 15 million single-cell images based on 16 publicly available image datasets. More than half of these images have annotated cell type labels. To ensure consistent annotations across the dataset, we aligned cell type names at varying levels of granularity, making them compatible with most existing annotations. We randomly partitioned this dataset into training and validation sets (25:1) to train the ensemble model.

We trained five models using those training images that contained the full set of markers for a given panel (Figure 1**C**). Each model was a customized ViT pre-trained and fine-tuned on `SPData_1` (model hyperparameters are listed in Table S2). Three are used to annotate immune cells, one for tissue structures and proliferating cells, and one for nerve cells. Note that CD45 in the tissue structure and nerve cell panels serves as a negatively expressed marker to distinguish immune cells. The accuracies of the immune cells models (blue bars in Figure 2**B**) on our held-out validation set demonstrate the effectiveness of these models. After applying all base models for which sufficient markers are present, final predictions for each cell are made by comparing prediction confidences from the base models. Moreover, for cells that are not annotated to known cell types, RIBCA supports grouping them into proposed additional cell types using unsupervised clustering approaches, as illustrated in Figure S2.

**Fig. 2.**
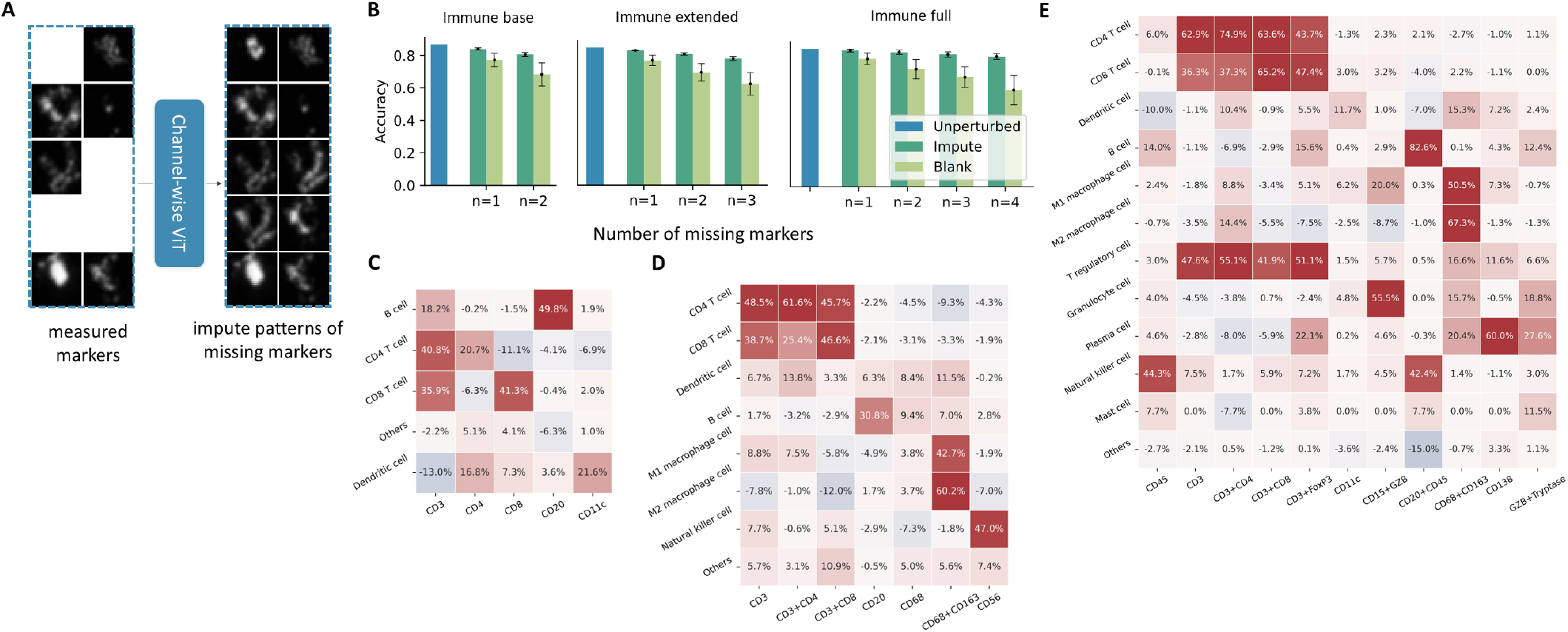
**A**, A ViT trained by channel-wise masked auto-encoding strategy can predict any missing image channels given the observed channels. **B**, Accuracies of immune panel models (trained using `SPData_1`) when evaluated with full images or averaged over removing one or more channels and either replacing it with a blank image or by imputation. The analysis presents accuracy differences (imputation method minus blank image) for various cell type models **C**, immune base, **D**, immune extended, and **E**, immune full models when the marker(s) (row) is missing and imputed or replaced by a blank image channel.

### RIBCA is flexible to diverse imaging panels of markers

To enhance robustness, we addressed scenarios where a query image might be missing only one or a few markers needed for a panel by developing a missing marker imputation strategy that works for a specific panel. This uses ViTs trained using a channel-wise masked auto-encoding (MAE) strategy, which takes the image channels of observed markers as input and predicts images for any missing markers (Figure 2**A**). We trained one such model for each of the three immune cell panels. Imputing missing markers consistently improved cell type classification accuracy compared to using blank image channels (Figure 2**B**), with greater benefits as more markers were missing. When imputing 4 missing markers of the immune full panel, the difference between using imputed images and real images is only 4.79%; however, this difference is 25.32% without imputation. The success in imputing missing markers makes RIBCA adaptable to various commonly used marker panels. The differences in accuracy for each cell type with and without our imputation method are shown in Figure 2**C**-**E**. It shows that even if critical markers are missing, the classifier can still have a good chance to predict the correct cell type. For example, when both CD3 and CD8 markers are missing, the accuracy of classifying a CD8^+^ T cell image is 0 without marker imputation but increases to 65.16% when using imputed images of CD3 and CD8 with the immune full panel model.

### RIBCA successfully annotates cell types in new experiments

To assess the generalizability of the ensemble on images from completely new experiments (and tissue types not present in `SPData_1`), we generated a dataset using the Human Protein Atlas (HPA) tissue microarrays (TMA) (referred as `SPHPAData_1`). In total, we generated 148 images of TMA cores from 46 different human tissues, acquired using a 46-plex panel (Table S3). This dataset is released in v24 of the HPA, and will be useful to the community for benchmarking purposes.

Examples of TMA multiplexed images and their annotations from RIBCA (without any fine-tuning) are shown in Figure 3**A** and **B**. The results align with biologists’ expectations in two key ways: (1) the spatial arrangement of cells, such as epithelial cells along intestinal crypts and B cells clustering in the spleen’s germinal center; (2) marker expression, evidenced by clear signals in the cell membranes of CD4 and CD8 T cells. To quantitatively assess performance, we manually validated selected images, achieving a high accuracy of approximately 82.2% in fine-grained cell type annotation. We also compared it with a pixel-based clustering method for cell type annotation, Pixie (7), that does not require training. For 2 out of the 7 TMAs we examined, Pixie was only able to assign 2-3 cell types, resulting in extremely low accuracies. For the remaining 5 TMAs, it achieved an overall accuracy of 62.34 %, 20 % lower than RIBCA. (RIBCA’s runtime for 7 TMA cores was also dramatically lower than Pixie’s (Figure 3**E**).) RIBCA generally predicts most cell types accurately, with the exception of stromal cells, which are often misclassified as epithelial cells. Additionally, the average marker expression across all TMA images (Figure 3**D**) and the correspondence between cell types and their signature markers meet expectations. We used RIBCA to efficiently annotate over 1 million cells in `SPHPAData_1`, revealing the spatially resolved cell type composition across different tissue samples, as shown in Figure 3**F**. The relative ratio of cells in the different tissues matched their previously reported and widely accepted functions. Tubular organs (e.g., GI tract) have a higher epithelial proportion than solid organs (liver and spleen). In the lymphatic system (lymph nodes, spleen, thymus), T and B cells dominate, while macrophages are more abundant in the liver, where Kupffer cells play a key role in maintaining homeostasis (8, 9).

**Fig. 3.**
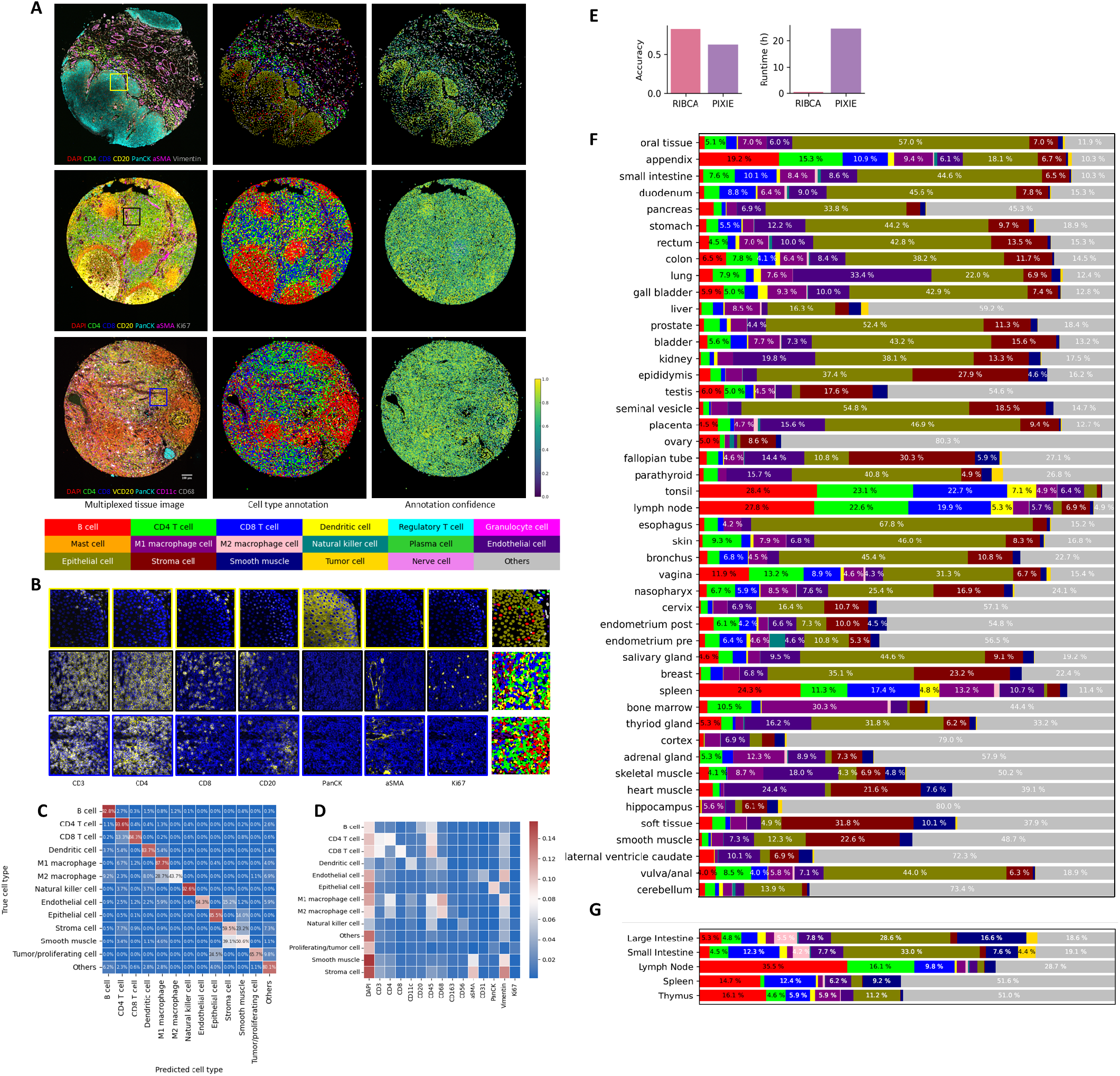
Results for diverse tissues. **A**, Images of DAPI and six selected markers or TMA cores (left) from the stomach, lymph node, and seminal vesicle (top to bottom). along with cell-level annotations (middle), and associated annotation confidence (right). **B**, Magnified regions (boxes in left column above) highlight cell nuclei (blue) and seven selected markers (yellow). The rightmost column shows the matching cell type annotations for this region using the same color scheme as above. **C**, Confusion matrix comparing cell type annotations by our model with manual annotation for selected TMA cores. **D**, Average protein expression for cell types annotated by the ensemble model across all TMA images. **E**, Comparison of cell type annotation accuracy validated on selected HPA TMA images between RIBCA and Pixie methods. Cell type distributions derived from our annotations across different tissue types for two datasets **F** `SPHPAData_1` and **G** HuBMAP CODEX images.

A set of 88 HuBMAP CODEX datasets from 5 tissues were also annotated. Those datasets varied somewhat in the markers present in each dataset, limiting which panels RIBCA could apply. The cell type results, along with which panels could be run, are shown in Figure S3. Figure 3**G** shows average results for datasets containing sufficient markers to allow annotation of at least the immune extended and the structure panels. Comparing the HPA and HuBMAP results, the cell type compositions are similar for small intestine and thymus, and small and large intestine are similar in the HuBMAP results. Those three tissues were analyzed by the Stanford Tissue Mapping Center. The images of the two other tissues were provided by the University of Florida Center and had only 11 markers, presumably accounting for the results with a large percentage of unidentified cells.

### RIBCA enables spatial analysis of cellular arrangement

RIBCA also enhances downstream analyses, such as tissue microenvironment and cell interaction studies. Figure 4**A** shows the average compositions of cell types within spatial niches (15 neighboring cells), facilitating the investigation of cell interaction heterogeneity. Specifically, we found that (1) dendritic cells have the most diverse neighbor composition, typically including various immune cells, reflecting their role as antigen-presenting cells crucial for immune cell differentiation and proliferation. B cells tend to cluster and form germinal centers in secondary lymphoid tissues, where they proliferate and activate. (2) Epithelial and stromal cells, as supporting tissues, are naturally located near each other, providing the structural foundation of the body. Highly proliferative epithelial cells are often interspersed among other epithelial cells or located in the epithelium’s uppermost layer. These neighborhood relationships highlight inherent cell functions. Furthermore, we sought to systematically partition tissue images into distinct regions using the RIBCA’s cell type annotations. This is realized by examining the cell type composition within the local cellular environment of each cell. Note that we intentionally do not use cell neighboring information when identifying cell types, this enables an unbiased partition in tissue regions. As illustrated in Figure 4 **B**, we employed the RIBCA’s annotated cell types (left) to identify distinctive tissue regions within lymph node and small intestine samples. In the lymph node, we identified four anatomical regions: the subcapsular sinus, paracortex, germinal center, and medulla. Each of these areas represents a specialized microenvironment with distinct cellular characteristics that contribute to the lymph node’s complex immunological functions. Similarly, our examination of the small intestine tissue uncovered four defined regions region: epithelium, submucosa, mucosa, and muscularis. These regions reflect the intricate structural and functional diversity inherent in intestinal tissue architecture. These findings highlight RIBCA’s potential to provide novel insights into the spatial organization of tissue functional units.

**Fig. 4.**
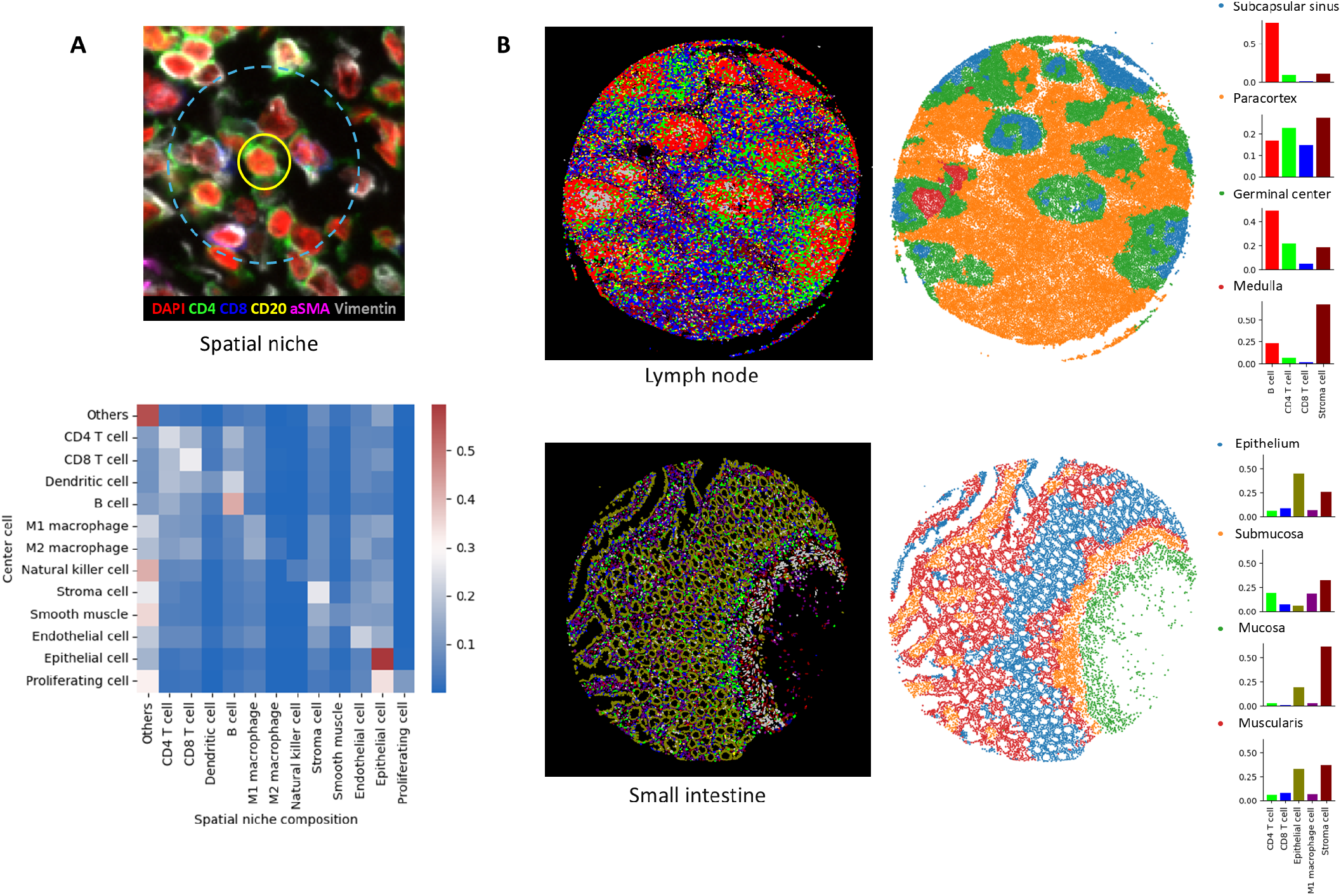
Results of RIBCA enabled spatial analysis. **A**: Neighboring cell composition for different cell types, with each row showing the normalized average fraction of neighboring cell types for the associated cell type. **B**: Examples of cell type annotation maps and corresponding tissue anatomical regions identified based on the cell type composition within the local cellular environment of each cell.

## Discussion

Exploring the frontier of spatial biology involves identifying cell types at the single-cell level in multiplexed proteomic images. Our model enables fast, unbiased, and automated cell type annotation for any imaging-based tissue proteomic data. It is particularly useful for exploration of complex spatial organization of different types of cells *in situ*. The models were empirically validated to be robust and generalizable to new images acquired by different assays. In general, RIBCA unlocks a major bottleneck in spatial proteomics, and the modular design allows the community to expand it to cover more cell types and states.

## Materials and Methods

### Dataset curation for ensemble model

We collected 15 datasets from various sources, as listed in Table S1 and Figure S4. For each dataset, we first performed sanity checks by randomly picking a few images and visually inspecting them to verify that they corresponded to the descriptions in the original papers. For published datasets without cell segmentation, we performed cell segmentation using CellPose 3.0. For datasets with annotated cell types but no cell segmentation masks, we did an additional step that maps our cell segmentation with the cell coordinates provided to ensure the correspondence between the images and cell type labels. For provided cell coordinates that did not match any of our segmented cells, we generated a circular pseudo-mask centered at that coordinate as a replacement for cell mask.

We also created a cell type marker list for each individual model (which we refer to as a panel) as shown in Table 1, and matched markers associated with each dataset to our panels accordingly. In cases where a marker was absent, we substituted it with a blank image, under the assumption that the absence of a specific marker indicates the corresponding cell type is not present in that dataset. To minimize these substitutions, in rare cases, we replaced some channels with closely related markers; for example, we used CD21 instead of CD20 and occasionally CD38 instead of CD138 in lymphoma datasets (10) for plasma cells. Although these markers serve distinct functional roles, they are considered interchangeable to some extent as cell type markers. Due to differences in granularity of cell type annotations and markers across different dataset, we aimed to maximize the use of the original labels. We manually assigned cell types for panels at different levels of granularity. The correspondence (or assignments) of original cell type annotations to our consolidated cell types (and panels) can be found in Tables S5-S9.

**Table 1.** Panel names, marker lists and corresponding cell types.

| Panel Name | Marker List | Cell Type names |
| --- | --- | --- |
| Basic Panel (Immune) | CD45, CD20 (or CD79a), CD4, CD8, DAPI, CD11c, CD3 | B cells, CD4 T cells, CD8 T cells, Dendritic Cell |
| Extended Panel (Immune) | DAPI, CD3, CD4, CD8, CD11c, CD20 (or CD79a), CD45, CD68, CD163, CD56 | CD4 T cell, CD8 T cell, Dendritic cell, B cell, Macrophage, M2 Macrophage, NK cell |
| Full Panel (Immune) | DAPI, CD3, CD4, CD8, CD11c, CD15, CD20 (or CD79a), CD45, CD56, CD68, CD138, CD163, FoxP3, Granzyme B, Tryptase | CD4 T cell, CD8 T cell, Dendritic cell, B cell, Macrophage, M2 Macrophage, Treg cell, Granulocyte (Neutrophil), Plasma cell, NK cell, Mast cell |
| Structure Panel | DAPI, aSMA, CD31, CK, Vimentin, Ki67, CD45 | Stroma, Smooth Muscle, Endothelial, Epithelial, Proliferating/Tumor |
| Nerve Panel | DAPI, CD45, GFAP (or Chromogranin A) | Nerve cell |

**Table 2.**
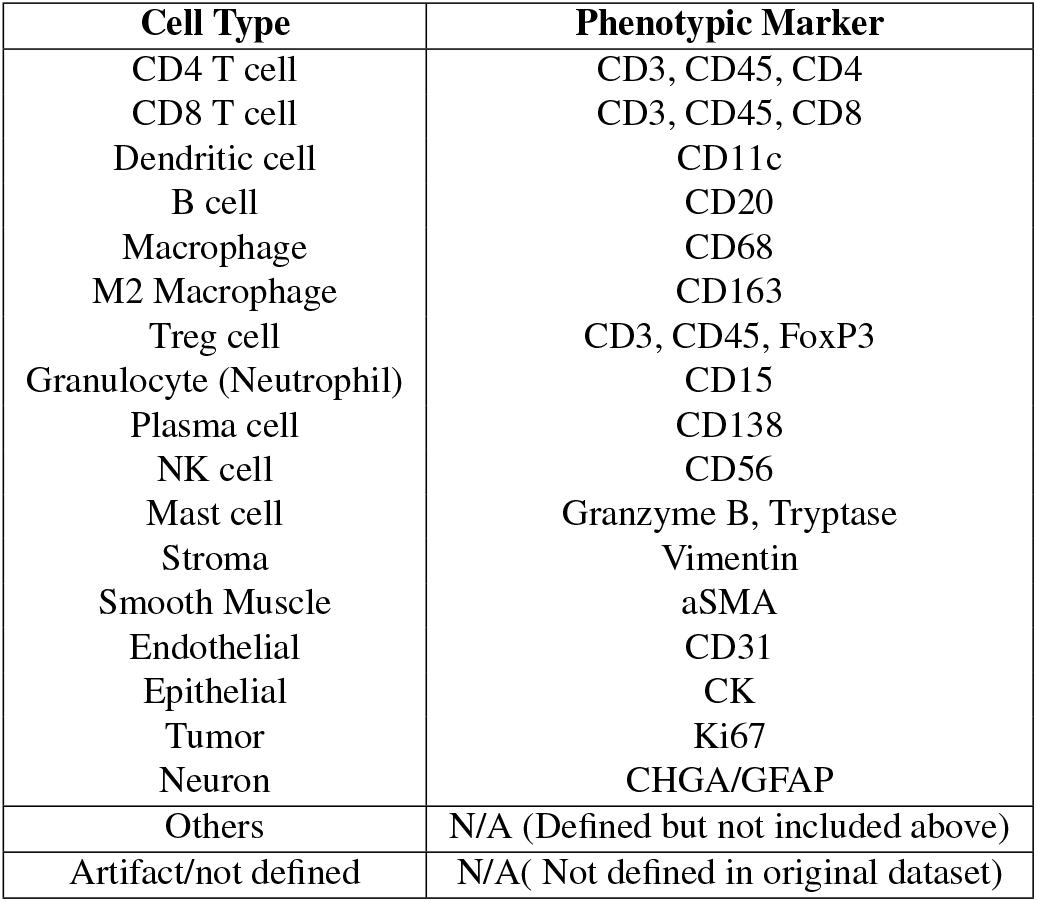
Phenotypic Markers and Their Corresponding Cell Types.

We used the unlabeled dataset for pretraining and the labeled dataset for fine-tuning. The datasets are summarized below. Scripts to regenerate the collection of datasets are provided (see Availability).

1. The Human colorectal cancer CODEX dataset consists of 140 tissue CODEX images from 35 advanced-stage colorectal cancer patients, using a panel of 56 proteins. This dataset consists of over 220,000 cells with 29 cell types. Cell types were identified through a process of manual gating, clustering, and assigning cluster groups to specific cell types. (11)
2. The Human Intestine CODEX dataset comprises 64 CODEX images of human colon tissue, with a panel of 47 proteins. It includes over 210,000 cells, with 21 distinct cell types. Cell type annotations were performed through clustering, followed by manual labeling of the clusters. (12)
3. The Lymphoma CODEX dataset includes 35 tumor-free lymphoid tissue cores from 19 patients, analyzed using a panel of 53 proteins. It comprises over 5.3 million cells, classified into 19 distinct cell types. Cell type annotation was performed using Leiden-based clustering on phenotypic markers, followed by merging of related clusters. (13)
4. The bone marrow and Acute Myeloid Leukemia (AML) CODEX dataset contains 7 AML and 12 bone marrow CODEX images, with a panel of 54 proteins. It includes over 1 million cells, with 37 distinct cell types. Cell type annotations were performed through Louvain clustering, manual annotation of clusters, and lateral spillover correction. (14)
5. The triple negative breast cancer MIBI-TOF dataset include 64 MIBI data from triple negative breast cancer, with a panel of 36 proteins. It includes over 60,000 cells, with 17 distinct cell types. Cell type annotations were performed by clustering in a hierarchical scheme, including FlowSOM of immune and non-immune cells and merging similar clusters by hierarchical clustering. (15)
6. The breast cancer progression MIBI dataset contains 79 MIBI samples from patients with ductal carcinoma in situ and invasive breast cancer, using a panel of 37 proteins. It includes over 200,000 cells. This dataset does not include annotations. (16)
7. The human tuberculosis MIBI-TOF dataset contains 17 MIBI samples from tuberculosis patients, using a panel of 29 proteins. It includes over 22,000 cells, representing 20 distinct cell types. Cell type annotations were performed using FlowSOM clustering. (17)
8. The fetally derived extravillous trophoblast MIBI dataset contains 211 images from the human maternal-fetal interface, utilizing a panel of 32 proteins. It includes over 400,000 cells, representing 25 distinct cell types. Cell type annotations were performed using a deep learning pipeline, supported by 93,000 manual annotations. (18)
9. The human kidney CODEX image dataset consists of 25 images from 56 healthy adult kidney tissues, using a panel of 21 proteins. It includes over 860,000 cells. This dataset does not have annotations. (19)
10. The HuBMAP 29-marker CODEX dataset contains 10 images from healthy adult spleen, thymus, and lymph node tissues. It includes over 940,000 cells. This dataset does not have annotations.
11. The classic Hodgkin Lymphoma CODEX dataset consists of one image from cHL patients, utilizing a panel of 49 proteins. It includes over 130,000 cells, representing 17 distinct cell types. Cell type annotations were conducted using traditional iterative clustering and visual inspection. (10)
12. The Tonsil CODEX Image I dataset consists of 45 images from healthy tonsil tissue, using a panel of 88 proteins. It includes over 110,000 cells. This dataset does not contain annotations.
13. The multi-tumor CODEX image dataset comprises 61 images from various tumor tissues, using a panel of 88 proteins. It includes over 240,000 cells. This dataset does not contain annotations.
14. The lung dataset comprises 3 images from human lung cancer tissue, using a panel of 27 proteins. It includes over 297,000 cells. This dataset does not contain annotations. (20)
15. The cHL CODEX dataset contains one FOV obtained using a 50-marker CODEX panel, comprising approximately 145,000 cells. An iterative approach with Rphenoannoy and FlowSOM (21, 22) categorized the majority of these cells into 16 distinct phenotypes, each averaging over 8,000 cells.
16. The Tonsil CODEX dataset II consists of one FOV image of human tonsil tissue, detected using a panel of 56 antibodies, as presented in (23). Over 120,000 cells were identified and grouped into 10 common cell types.

### Cell Type Annotation Benchmarking Comparison

We compared results from the ViT model with those from three other recently published representative methods, CellSighter (3) and MAPS (10), and one classic method, STELLAR (24), by testing them on 5 public datasets with their provided software. The five data sets are the cHL MIBI, 4 FOVs of human small intestine (including over 150,000 cells, as used for benchmarking in (24)), cHL CODEX, tonsil CODEX datasets, and one additional Melanoma Lymph Node dataset that was not included in `SPData_1` (published in (3)). This dataset includes 16 FOVs of MIBI images from lymph nodes affected by melanoma metastases across multiple patients, using a panel of 25 protein markers. Around 115,000 cells were identified through a process involving FlowSOM, gating, and visual inspection, followed by manual annotation. This approach led to the identification of 15 distinct cell types.

For each dataset, we randomly partitioned it into a small training dataset (10% of total data) and a validation set (90%), where each cell image/profile in the training set is accompanied by their associated cell type annotation. We independently repeated all tests 7 times and reported the average scores.

### Full Field of View Image Preprocessing

Our method applies to registered and stitched 2-dimensional multiplexed images. (it can be extended for 3-dimensional images in general.) First, we subtracted the background signal using the rolling-ball method for each image channel individually (25). Next, for each image channel, the single-channel intensity was clipped at the 98th percentile of nonzero pixels, i.e.,

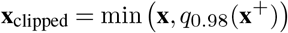

given **x** denotes the (flattened) image vector, **x**^+^ denotes the set of nonzero elements in **x** and *q*_0.98_(·) as the 98th percentile. Then, the clipped single-channel image was linearly scaled into the range of -1 to 1, i.e.

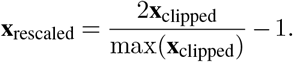

Finally, for images acquired using multiplexed ion beam imaging (MIBI) method, we applied Gaussian blurring with *σ* = 0.5 to each channel of the image.

Our approach assumes cell-level segmentation associated with the multiplexed images in use. Segmentation can be obtained from tools like Cellpose (26, 27) and Deepcell Mesmer (28). In this work, we used the cell segmentation provided with the datasets or Cellpose 3.0 to segment cells when the segmentation was not provided.

### Soft-masking of Individual Cell Images

For each segmented cell, a square image tile with a side length of 40 pixels centered on the cell was cropped out of the full image. However, since the resulting single-cell image may still contain protein patterns belonging to its neighboring cells, we applied a *soft-masking* operation; the softened cell mask is essentially an average of original cell mask being morphologically dilated and Gaussian blurred using different parameters. That is, for a cell image, we denote its associated cell mask (from the original segmentation) as **b** ∈ {0, 1} ^40×40^, and refer `dilate`(·, *r*) to the binary dilation operation of an image using a disk-shaped kernel, with *r* specifying the radius, and `blur`(·, *r*) as the Gaussian blur operation with kernel radius *r*. Then, a soft-mask is written as 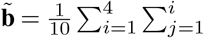 `blur`(`dilate`(**b**, *j*), *i*). To apply softmasking to individual images, we simply perform elementwise multiplication between 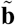 and each image channel. Intuitively, soft-masking produces a smoother transition from the inside to outside of the cell region, avoiding sudden changes of intensity on the edges of the cell region. An example of soft-masking of an individual cell region is shown in Figure S5

### Vision Transformer

ViT is a widely used neural network design introduced by Dosovitskiy *et al*. (6). We briefly review its mechanism here. The idea of ViT is to partition an image into a sequence of non-overlapping patches, and use a Transformer encoder neural network originally designed for 1-dimensional sequences to transform this array of image patches. In particular, given an input image **x** ∈ ℝ^*C*·*H*·*W*^ , it will be cropped (on the plane of *H* × *W* ) into a grid of patches in size *C* · *P* · *P* , resulting in 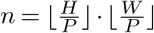 patches in total denoting as 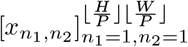. Each patch 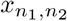 is then flattened as a 1-dimensional vector and linearly transformed to dimensionality of *d, i*.*e*. 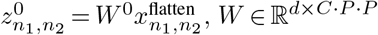 . These transformed vectors are also called tokens of this image. Following the original design of ViT, a `class` token (a learnable *L*-dimensional vector) is prepended to this sequence of tokens, 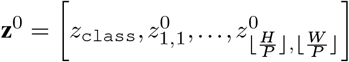 , the intuition of this operation is to use this additional vector as a summarized feature of the rest of transformed patch vectors after ViT encoding. To preserve the spatial arrangement of the patches in the original image, another sequence of *d*-dimensional vectors generated by periodic functions is added to **z**^0^ to indicate the position of each patches in the original image, realized by **z**^1^ = **z**^0^ + **P, P** ∈ ℝ^*n*+1×*d*^,

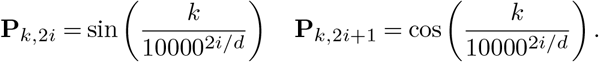

Next, these positional-aware tokens are further encoded by a multi-head self-attention based Transformer encoder neural network as described (29). We denote the encoded tokens as **z**^enc^ = `enc`(**z**^1^)∈ ℝ^*n*+1×*d*^. To map the encoded tokens onto the classification decision, a linear map is applied to the encoded `class` token, *i*.*e*. the first element of **z**^enc^, say 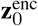 and 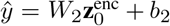 where *W*_2_ ∈ ℝ^*m*×*d*^ and *b* ∈ ℝ^*m*^, *m* is the number of classes and ŷ_*u*_*n* denotes the unnormalized probabilities.

### Masked Auto-encoding

Dosovitskiy *et al*. (6) showed that unsupervised pre-training of a ViT encoder can improve its performance of image classification. A popular pre-training method is masked auto-encoding MAE. MAE aims to recover the original full image from a partial observation of that image. The realization of MAE relies on a ViT as an image encoder. Instead of receiving all patches cropped from the input image, during MAE pre-training, only a small subset of patches are *observed* and used as the input to the ViT encoder (and no `class` prepended). To decode the image from tokens, the encoded tokens associated with the observed patches are placed in their original position in the image, and the positions associated with unobserved patches are filled by a vector (the same size as each encoded token) called mask representation (a model parameter used as the representations of unobserved patches), this sequence of tokens are then decoded by a simpler decoder model. That is, only *m* = ⌊(1 −mask ratio)*n*⌋ patches are observed, denoting their indices as *I*. They are linearly transformed to *d*-dimensionality, added with their associated positional tokens, and encoded by the ViT encoder as,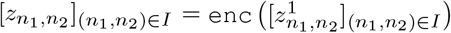 , then the mask representation will be inserted, i.e. 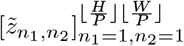 where 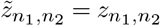 if (*n*_1_ , *n*_2_ ) ∈ *I*, otherwise,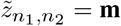. Here, **m** denotes the token associated with any masked patches. A decoder will decode the these tokens to reconstruct the original image, as 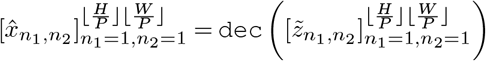 , and the mean square error (MSE) is calculated between the original signal and this reconstruction. The intuition of MAE is to use a simple but effective way to learn spatial relationships between different parts of images without any references and pre-train a scalable visual representation learning model. After MAE pre-training, only the pre-trained ViT encoder is preserved.

To apply it for cell type identification tasks, we pre-trained our ViT encoder on images cropped from a full-view multiplexed tissue image, each containing only one single centered cell with very small margins around. In practice, we encoded each individual cell image (as described in Methods Section) using the ViT encoder, as illustrated in Figure 1**d**. We used a small ViT as encoder; the hyperparameters of our models are listed in Table S2. The models were trained to minimize the average mean square error of each mini-batch using an Adam optimizer with an adaptive learning rate of 10^−3^.

To train a channel-wise ViT using the MAE approach, we first split a multi-channel image into individual image channels, and tile them into a larger single-channel image. For example, given an image with the size of *C*× *H*× *W* and *C* = *ab, a, b* are integers, we can tile it into a new image with a size of 1 × *aH* × *bW* . By choosing the patch size equal to 1 × *H* × *W* which results in *C* patches in total, we can train a ViT using the MAE technique described in this section by randomly masking patches in this larger image (that is, individual image channel of the original image). In particular, we did not use a fixed number of masked patches (image channels) during this training, rather we randomly masked nearly half (plus or minus two) of the patches to train this model.

### Fine-tuning

After pre-training, we fine-tuned the ViT model following a standard classification task training paradigm. The weights of the ViT encoder parameter are borrowed from the pre-trained model, and the rest of parameters are initialized randomly according to a Gaussian distribution (standard deviation of 0.02). Given some reference cell type annotation, say 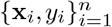 provided **x**_*i*_, *y*_*i*_, *i* ∈ [*n*] are singlecell images and their associated cell type labels, the ViT encoder encodes **x**_*i*_ into tokens. Next, the encoded `class` token is mapped onto unnormalized probabilities *ŷ*_*u*_*n* as described previously, which are further normalized by a Sigmoid function. The normalized predicted probability is used to calculate the label smoothing cross-entropy loss (30), as

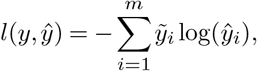

where 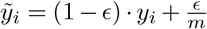 and *m* is the number of classes. Whole ViT model is trained by an AdamW optimizer with a learning rate of 2 × 10^−4^.

### Merging predictions

As the ensemble model possesses multiple individual models, a necessary step before final classification is to merge the decisions from applied models. For the three immune panels, the larger model is prioritized. For example, when both extended and base models can be used, only the extended model is applied. To merge the predictions across other panels, the predicted normalized probabilities (scores) from all relevant models are collected. To establish a cutoff for determining whether a specific cell type should be assigned, we take the lower of (1) the minimum of all scores for the “Others” class and (2) a user-specified hyperparameter (set to 0.3 for the analyses here). If the cell type with the highest score (other than the class of “Others”) is greater than this value, it is assigned; othewise it is assigned “Others”. All cells assigned to “Others” can be further grouped into additional cell types using classic clustering methods such as HDBSCAN method (**?** ) provided the minimal number of cells that form a cell type.

### Runtime

For a single TMA core image with 10,000 cells, it takes about 5 to 10 minutes to generate cell type annotation maps and all downstream analyses on an ordinary desktop computer.

### Expert Validation of the Ensemble Model

To validate the assigned cell types, we chose a small region of interest in each TMA and checked their corresponding marker signals. To ensure the composition of different cell types, we chose the region based on the criteria that the area contains different compartments. Based on the spatial organization of markers, certain tissues appear to have a clear compartmentalization. These compartments were visually determined by the enrichment of markers. As an example, for intestine tissue, at least two layers should be included, which would include both epithelial and immune cells. Epithelial layers are enriched in cytokeratin expression and immune cells layer are enriched in CD45 signals, with a clear boundary between them.We want to ensure that the model is able to distinguish between different tissue compartments at the boundary where the enrichment of cells is different, and label assignment is not affected by signal spillover. For each selected region, we manually validated the cell assignments using the Napari plugin we developed. This plugin allows us to visualize both RIBCA’s annotation and multiplexed data together. For each cell, we validated its annotation by checking if its signature markers were fully expressed. Cells with valid marker expression were considered correctly classified. When discrepancies were observed, we manually reviewed the phenotypic markers to identify the positive ones and reassign the cell to the correct type. We repeated this procedure for each selected cell to plot the confusion matrix as shown in Figure 3**C**.

### Tissue region identification

Inspired by (**?** ) who used first-order spatial statistic for tissue region partitioning, we developed a computationally simpler method that identifies distinct tissue functional regions and analyzes the cellular neighborhood composition around each cell. Our approach began with RIBCA’s cell type annotation, after which we characterized the spatial context of each cell by examining its local cellular environment. Specifically, we calculated the cell type fractions within progressively expanding neighborhoods, ranging from the 10 nearest cells to the 200 (10, 20, 30, 50, 75, 100, 150, 200) nearest cells. For each cell, we created a spatial feature vector by concatenating the cell type fractions derived from these different neighborhood sizes. That is, denoting *C* as the total number of cell types, we write 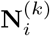 as the collection of *k* nearest cells of cell *i*, and denoting **CT**(·) as a function that returns the cell type of the input cell instance, the spatial feature vector for cell *i* is written as

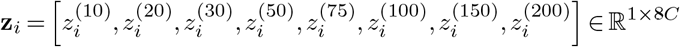

where 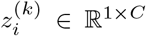, and the *j*-th entry of 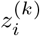 is 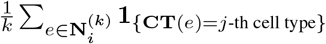.This approach allowed us to capture the intricate spatial relationships between different cell types at multiple scales. Once we generated these spatial features for every cell in the image, we applied spectral clustering (**?** ) to segment the tissue into distinct regions. Each resulting cluster represents a tissue region characterized by its unique cellular composition and spatial arrangement.

### Human tissue sample preparation

We followed the tissue sample preparation protocol described by Danielsson *et al*. (31). Tissue samples were handled in accordance with Swedish laws and regulation and obtained from the Department of Pathology, Uppsala University Hospital, Sweden, as part of the sample collection governed by the Uppsala Biobank (http://www.uppsalabiobank.uu.se/en/). All human tissue samples used were anonymized in accordance with approval and advisory report from the Uppsala Ethical Review Board (Dnr Ups 02-577). TMAs were generated in accordance with strategies used in the Human Protein Atlas and as previously described (32, 33). In brief, hematoxylin-eosin stained tissue sections from formalin-fixed paraffin-embedded donor blocks were examined in order to verify the histology and select representative regions for sampling into the TMA. Normal tissue was defined as microscopically normal and was most often selected from specimens collected from the vicinity of surgically removed tumors. Two TMA blocks were generated containing triplicate 1mm cores of 45 total types of normal tissue (tissues listed in Table S4).

### Spatial targeted proteomic profiling using PhenoCycler Fusion platform

Highly multiplexed protein data were generated using the PhenoCycler Fusion dual-slide system (Akoya Biosciences, Marlborough, MA). The staining was performed following the PhenoCycler-Fusion User Guide_2.1.0 version. We cut a 4 µm thick section of TMA block in a positively charged micro-scope slide (VWR, Epredia, Cat No. 76406-502) and bakedit at 65 °C in an oven, overnight. The next day, we manually dewaxed the sections starting with immersion in HistoChoice (Sigma, Cat No. H2779), 5 minutes twice, and rehydrated in descending concentrations of ethanol (100% ethanol twice, 90%, 70%, 50%, 30%, ddH_2_O twice; each step for 5 minutes). We performed then heat-induced epitope retrieval (HIER) treatment using pH 9 EDTA buffer (Akoya Biosciences, Cat No. AR900250ML) in a pressure cooker (Bio SB TintoRetriever, Cat No. BSB-7087) at 114–121 °C during 20 minutes. Once the slides were equilibrated at room temperature, we transferred them to a ddH_2_O container for a total of two washes, each step of 2 minutes. To reduce the tissue autofluroescence, we performed a photobleaching treatment: we submerged the slides in a transparent slide mailer (Heathrow Scientific, Cat No. HS15982) containing 25 ml 1×PBS (Gibco, Cat No. 18912-014), 4.5 ml 30% H_2_O_2_ (Sigma, Cat No. 216763) and 0.8 ml 1M NaOH (Sigma, Cat No. S5881). The slides in solution were exposed to two LED lamps (GLIME, Light Therapy Lamp UV Free 32000), one on each side, and incubated for 45 minutes. We prepared a new photobleaching solution and incubated it for an additional 45 min, following a total of four 1×PBS washes, each step of 2 minutes. The slides were transferred to a Coplin Jar containing Hydration Buffer (Akoya Biosciences, Cat No. 7000017) and incubated for 2 minutes, twice; we moved the slides to a jar with Staining Buffer (Akoya Biosciences, Cat No. 7000017) and equilibrated for 30 minutes. Meantime, we diluted all the antibodies (each antibody was conjugated with an unique oligosequence, antibody dilutions listed in Table S3) together in a cocktail solution containing Staining Buffer and N, J, G and S blockers (Akoya Biosciences, Cat No. 7000017) and added them on the slides for an overnight incubation at 4 °C. The day after, the primary antibodies were removed from the slides through washes on Staining Buffer and then we performed 3 different consecutive post-fixation steps with washes between them: 1) 1.6% paraformaldehyde (ThermoFisher, Cat No. 043368) diluted in Storage Buffer (Akoya Biosciences, Cat No. 7000017) for a 10 minutes incubation at room temperature; 2) ice cold methanol (Sigma, Cat No. 322415) for 5 minutes at 4 °C; 3) Fixative Solution (Akoya Biosciences, Cat No. 7000017) diluted at 1:50 in 1×PBS. The slides were stored in the Storage Buffer till they were attached to the respective flow cells (Akoya Biosciences, Cat No. 7000017) to be moved to the PhenoCycler Fusion system to start the experiments. In addition, we prepared two 96-well reporter plates that contained a fluorescent dye conjugated to a PhenoCycler oligonucleotide sequence complementary to one specific antibody barcode: unique reporters were added by groups of up to 3 spectrally different dyes plus nuclear stain in each single well.

### PhenoCycler image acquisition

We performed the image acquisition on the PhenoImager Fusion 2.2.0 (Akoya Biosciences, Marlborough, MA) where three fluorescent oligo reporters with spectrally distinct dyes were applied to the tissue in iterative imaging cycles. The images were captured using a widefield microscope with a Olympus UCPlanFL 20×/0.70 NA objective and a resolution of 0.5 µm/pixel. Whole-slide images were automatically aligned using DAPI as a reference, and background subtracted to be assembled into the the final QPTIFF image files.

## Availability

RIBCA is available at https://github.com/sun-huangqingbo/multiplexed-image-annotator. Since the size of the full `SPData_1` dataset prevents its inclusion in the repository, we include scripts (https://github.com/sun-huangqingbo/SPData_1-curation to reproduce the processing of the original dataset images to our cropped cell images. The raw image sources are listed in Table S1.

The HPA tissue microarray multiplexed images are available in the Human Protein Atlas (v24).

## ACKNOWLEDGEMENTS

This work was supported in part by the Wallenberg Foundation (2021.0346) and as part of the PROMINENT team supported by the Cancer Grand Challenges partnership funded by Cancer Research UK (CGCATF-2021/100010), the National Cancer Institute (OT2CA278713), and the Scientific Foundation of the Spanish Association Against Cancer, AECC, and by grant OT2 OD033761 from the National Institutes of Health Common Fund.

## Bibliography

1. Hongkui Zeng. What is a cell type and how to define it? Cell, 185(15):2739–2755, 2022.

2. Weiruo Zhang, Irene Li, Nathan E Reticker-Flynn, Zinaida Good, Serena Chang, Nikolay Samusik, Saumyaa Saumyaa, Yuanyuan Li, Xin Zhou, Rachel Liang, et al. Identification of cell types in multiplexed in situ images by combining protein expression and spatial information using celesta. Nature Methods, 19(6):759–769, 2022.

3. Yael Amitay, Yuval Bussi, Ben Feinstein, Shai Bagon, Idan Milo, and Leeat Keren. Cell-sighter: a neural network to classify cells in highly multiplexed images. Nature communications, 14(1):4302, 2023.

4. Ajit Johnson Nirmal, Clarence Yapp, Sandro Santagata, and Peter Sorger. Cell spotter (cspot): A machine-learning approach to automated cell spotting and quantification of highly multiplexed tissue images. bioRxiv, pages 2023–11, 2023.

5. Lorenz Rumberger, Noah F Greenwald, Jolene Ranek, Potchara Boonrat, Cameron Walker, Jannik Franzen, Sricharan Varra, Alex Kong, Cameron Sowers, Candace C Liu, et al. Automated classification of cellular expression in multiplexed imaging data with nimbus. bioRxiv, pages 2024–06, 2024.

6. Alexey Dosovitskiy, Lucas Beyer, Alexander Kolesnikov, Dirk Weissenborn, Xiaohua Zhai, Thomas Unterthiner, Mostafa Dehghani, Matthias Minderer, Georg Heigold, Sylvain Gelly, et al. An image is worth 16×16 words: Transformers for image recognition at scale. arXiv preprint 2010.11929, 2020.

7. Candace C Liu, Noah F Greenwald, Alex Kong, Erin F McCaffrey, Ke Xuan Leow, Dunja Mrdjen, Bryan J Cannon, Josef Lorenz Rumberger, Sricharan Reddy Varra, and Michael Angelo. Robust phenotyping of highly multiplexed tissue imaging data using pixel-level clustering. Nature Communications, 14(1):4618, 2023.

8. Oliver Krenkel and Frank Tacke. Liver macrophages in tissue homeostasis and disease. Nature Reviews Immunology, 17(5):306–321, 2017.

9. Ron Sender, Yarden Weiss, Yoav Navon, Idan Milo, Nofar Azulay, Leeat Keren, Shai Fuchs, Danny Ben-Zvi, Elad Noor, and Ron Milo. The total mass, number, and distribution of immune cells in the human body. Proceedings of the National Academy of Sciences, 120 (44):e2308511120, 2023.

10. Muhammad Shaban, Yunhao Bai, Huaying Qiu, Shulin Mao, Jason Yeung, Yao Yu Yeo, Vignesh Shanmugam, Han Chen, Bokai Zhu, Jason L Weirather, et al. Maps: Pathologist-level cell type annotation from tissue images through machine learning. Nature Communications, 15(1):28, 2024.

11. Christian M Schürch, Salil S Bhate, Graham L Barlow, Darci J Phillips, Luca Noti, Inti Zlobec, Pauline Chu, Sarah Black, Janos Demeter, David R McIlwain, et al. Coordinated cellular neighborhoods orchestrate antitumoral immunity at the colorectal cancer invasive front. Cell, 182(5):1341–1359, 2020.

12. John W Hickey, Winston R Becker, Stephanie A Nevins, Aaron Horning, Almudena Espin Perez, Chenchen Zhu, Bokai Zhu, Bei Wei, Roxanne Chiu, Derek C Chen, et al. Organization of the human intestine at single-cell resolution. Nature, 619(7970):572–584, 2023.

13. Tobias Roider, Marc A Baertsch, Donnacha Fitzgerald, Harald Voehringer, Berit J Brinkmann, Felix Czernilofsky, Mareike Knoll, Laura Llaó-Cid, Anna Mathioudaki, Bianca Faßbender, et al. Multimodal and spatially resolved profiling identifies distinct patterns of t cell infiltration in nodal b cell lymphoma entities. Nature Cell Biology, pages 1–12, 2024.

14. Shovik Bandyopadhyay, Michael P Duffy, Kyung Jin Ahn, Jonathan H Sussman, Minxing Pang, David Smith, Gwendolyn Duncan, Iris Zhang, Jeffrey Huang, Yulieh Lin, et al. Mapping the cellular biogeography of human bone marrow niches using single-cell transcriptomics and proteomic imaging. Cell, 187(12):3120–3140, 2024.

15. Leeat Keren, Marc Bosse, Diana Marquez, Roshan Angoshtari, Samir Jain, Sushama Varma, Soo-Ryum Yang, Allison Kurian, David Van Valen, Robert West, et al. A structured tumor-immune microenvironment in triple negative breast cancer revealed by multiplexed ion beam imaging. Cell, 174(6):1373–1387, 2018.

16. Tyler Risom, David R Glass, Inna Averbukh, Candace C Liu, Alex Baranski, Adam Kagel, Erin F McCaffrey, Noah F Greenwald, Belén Rivero-Gutiérrez, Siri H Strand, et al. Transition to invasive breast cancer is associated with progressive changes in the structure and composition of tumor stroma. Cell, 185(2):299–310, 2022.

17. Erin F McCaffrey, Michele Donato, Leeat Keren, Zhenghao Chen, Alea Delmastro, Megan B Fitzpatrick, Sanjana Gupta, Noah F Greenwald, Alex Baranski, William Graf, et al. The immunoregulatory landscape of human tuberculosis granulomas. Nature immunology, 23 (2):318–329, 2022.

18. Shirley Greenbaum, Inna Averbukh, Erin Soon, Gabrielle Rizzuto, Alex Baranski, Noah F Greenwald, Adam Kagel, Marc Bosse, Eleni G Jaswa, Zumana Khair, et al. A spatially resolved timeline of the human maternal–fetal interface. Nature, 619(7970):595–605, 2023.

19. Jens Hansen, Rachel Sealfon, Rajasree Menon, Michael T Eadon, Blue B Lake, Becky Steck, Kavya Anjani, Samir Parikh, Tara K Sigdel, Guanshi Zhang, et al. A reference tissue atlas for the human kidney. Science advances, 8(23):eabn4965, 2022.

20. Rumana Rashid, Giorgio Gaglia, Yu-An Chen, Jia-Ren Lin, Ziming Du, Zoltan Maliga, Denis Schapiro, Clarence Yapp, Jeremy Muhlich, Artem Sokolov, et al. Highly multiplexed immunofluorescence images and single-cell data of immune markers in tonsil and lung cancer. Scientific data, 6(1):323, 2019.

21. Jacob H Levine, Erin F Simonds, Sean C Bendall, Kara L Davis, D Amir El-ad, Michelle D Tadmor, Oren Litvin, Harris G Fienberg, Astraea Jager, Eli R Zunder, et al. Data-driven phenotypic dissection of aml reveals progenitor-like cells that correlate with prognosis. Cell, 162(1):184–197, 2015.

22. Sofie Van Gassen, Britt Callebaut, Mary J Van Helden, Bart N Lambrecht, Piet Demeester, Tom Dhaene, and Yvan Saeys. Flowsom: Using self-organizing maps for visualization and interpretation of cytometry data. Cytometry Part A, 87(7):636–645, 2015.

23. Sarah Black, Darci Phillips, John W Hickey, Julia Kennedy-Darling, Vishal G Venkataraaman, Nikolay Samusik, Yury Goltsev, Christian M Schürch, and Garry P Nolan. Codex multiplexed tissue imaging with dna-conjugated antibodies. Nature protocols, 16(8):3802– 3835, 2021.

24. Maria Brbić, Kaidi Cao, John W Hickey, Yuqi Tan, Michael P Snyder, Garry P Nolan, and Jure Leskovec. Annotation of spatially resolved single-cell data with stellar. Nature Methods, 19(11):1411–1418, 2022.

25. Stanley R Sternberg. Biomedical image processing. Computer, 16(01):22–34, 1983.

26. Carsen Stringer, Tim Wang, Michalis Michaelos, and Marius Pachitariu. Cellpose: a generalist algorithm for cellular segmentation. Nature methods, 18(1):100–106, 2021.

27. Marius Pachitariu and Carsen Stringer. Cellpose 2.0: how to train your own model. Nature methods, 19(12):1634–1641, 2022.

28. Noah F Greenwald, Geneva Miller, Erick Moen, Alex Kong, Adam Kagel, Thomas Dougherty, Christine Camacho Fullaway, Brianna J McIntosh, Ke Xuan Leow, Morgan Sarah Schwartz, et al. Whole-cell segmentation of tissue images with human-level performance using large-scale data annotation and deep learning. Nature biotechnology, 40(4):555–565, 2022.

29. Ashish Vaswani, Noam Shazeer, Niki Parmar, Jakob Uszkoreit, Llion Jones, Aidan N Gomez, Łukasz Kaiser, and Illia Polosukhin. Attention is all you need. Advances in neural information processing systems, 30, 2017.

30. Christian Szegedy, Vincent Vanhoucke, Sergey Ioffe, Jon Shlens, and Zbigniew Wojna. Rethinking the inception architecture for computer vision. In Proceedings of the IEEE conference on computer vision and pattern recognition, pages 2818–2826, 2016.

31. Angelika Danielsson, Fredrik Pontén, Linn Fagerberg, Björn M Hallström, Jochen M Schwenk, Mathias Uhlén, Olle Korsgren, and Cecilia Lindskog. The human pancreas proteome defined by transcriptomics and antibody-based profiling. PLoS One, 9(12):e115421, 2014.

32. Mathias Uhlen, Per Oksvold, Linn Fagerberg, Emma Lundberg, Kalle Jonasson, Mattias Forsberg, Martin Zwahlen, Caroline Kampf, Kenneth Wester, Sophia Hober, et al. Towards a knowledge-based human protein atlas. Nature biotechnology, 28(12):1248–1250, 2010.

33. Caroline Kampf, IngMarie Olsson, Urban Ryberg, Evelina Sjöstedt, and Fredrik Pontén. Production of tissue microarrays, immunohistochemistry staining and digitalization within the human protein atlas. JoVE (Journal of Visualized Experiments), (63):e3620, 2012.

