## Supplementary Figures and Tables for "Flexible and robust cell type annotation for highly multiplexed tissue images"

December 17, 2024

#### 1 Supplementary tables

Table S1: Sources of datasets used to curate our integrated dataset for training the models.

| Deposited data | Source Article | |
|---|---|---|
| Human colorectal cancer CODEX image | [10] | Link |
| Tonsil CODEX image I | | Link |
| Multi-tumor CODEX image | | Link |
| Lymphoma CODEX image | [9] | Link |
| Human intestine CODEX image | [5] | Link<br>Link |
| Bone marrow and acute myeloid leukemia CODEX image | [1] | Link |
| Tonsil CODEX image II | [2] | Link |
| Classic Hodgkin Lymphoma MIBI<br>Classic Hodgkin Lymphoma CODEX image | [11] | Link |
| Triple negative breast cancer MIBI-TOF | [6] | Link |
| Breast cancer progression MIBI-TOF | [8] | Link |
| Human tuberculosis MIBI-TOF | [7] | Link |
| Fetally derived extravillous trophoblasts MIBI | [3] | Link |
| HuBMAP 29-marker CODEX image (spleen, thymus, and lymph node) | | Link |
| Human kidney CODEX image | [4] | Link |

Table S2: Hyper-parameter setting of our customized ViT model.

| | ViT encoder | | | | Decoder | | | |
|---|---|---|---|---|---|---|---|---|
| Panel | Layer | Hidden size | MLP size | Heads | Layer | Hidden size | MLP size | Heads |
| Immune Base | 12 | 288 | 1152 | 12 | 6 | 192 | 768 | 8 |
| Immune Extended | 12 | 384 | 1536 | 12 | 6 | 192 | 768 | 8 |
| Immune Full | 12 | 576 | 2304 | 12 | 6 | 192 | 768 | 8 |
| Tissue Structure | 12 | 288 | 1152 | 12 | 6 | 192 | 768 | 8 |
| Nerve Cell | 12 | 144 | 576 | 12 | 4 | 96 | 384 | 4 |

Table S3: Antibody list multiplexed in each TMA section: all the antibodies were diluted at 1:200 dilution, except CD163 (1:50) and VIM (1:400). The exposure times were set up at 150ms for all the antibodies, except VIM (100ms). Akoya Biosciences is our antibody vendor.

| Target | Barcode | Fluorophore | Cat# |
|---|---|---|---|
| b-Actin | BX117 | AF750 | 4450092 |
| ATM | BX083 | Atto550 | 4250095 |
| Bcl2 | BX085 | AF647 | 4550089 |
| b-Catenin | BX096 | Atto550 | 4250091 |
| Caveolin | BX086 | AF647 | 4550084 |
| CD3e | BX045 | AF647 | 4550119 |
| CD4 | BX003 | AF647 | 4550112 |
| CD8 | BX026 | Atto550 | 4250012 |
| CD11c | BX024 | AF647 | 4550114 |
| CD14 | BX037 | Atto550 | 4450047 |
| CD20 | BX007 | AF750 | 4450018 |
| CD31 | BX001 | AF750 | 4450017 |
| CD34 | BX025 | Atto550 | 4250057 |
| CD39 | BX099 | Atto550 | 4250076 |
| CD40 | BX010 | AF647 | 4550064 |
| CD45 | BX021 | AF647 | 4550121 |
| CD44 | BX005 | Atto550 | 4450041 |
| CD56 | BX028 | AF647 | 4550126 |
| CD68 | BX015 | AF647 | 4550113 |
| CD107a | BX006 | AF647 | 4550098 |
| CD163 | BX069 | Atto550 | 4250079 |
| COL4A1 | BX042 | AF647 | 4550122 |
| E-cadherin | BX014 | Atto550 | 4250021 |
| EpCAM | BX091 | AF647 | 4550088 |
| ER (Estrogen receptor) | BX084 | Atto550 | 4250074 |
| GP100 (PMEL) | BX094 | AF647 | 4550091 |
| HIF1a | BX062 | AF647 | 4550069 |
| HistoneH3Phospho | BX030 | AF647 | 4550115 |
| HLA-A | BX029 | Atto550 | 4250100 |
| HLADR | BX033 | AF647 | 4550118 |
| ICOS | BX054 | AF647 | 4550117 |
| IDO1 | BX027 | AF647 | 4550123 |
| iNOS | BX023 | Atto550 | 4250073 |
| Keratin5 | BX101 | Atto550 | 4250090 |
| Keratin8/18 | BX081 | AF750 | 4450082 |
| Ki67 | BX047 | AF750 | 4450096 |
| Pancytokeratin | BX019 | AF750 | 4450020 |
| PCNA | BX036 | AF647 | 4550124 |
| PD1 | BX046 | AF647 | 4550038 |
| Podoplanin | BX121 | Atto550 | 4250094 |
| SMA | BX013 | AF750 | 4450049 |
| SOX2 | BX102 | Atto550 | 4250075 |
| TOX | BX060 | Atto550 | 4250067 |
| Vimentin | BX022 | AF750 | 4450050 |
| VISTA | BX040 | Atto550 | 4250063 |

Table S4: Tissue included in the TMA blocks to perform the highly multiplexed staining.

| | | | | |
|---|---|---|---|---|
| Adrenal gland | Cortex | Hippocampus | Parathyroid | Soft tissue |
| Appendix | Duodenum | Kidney | Placenta | Skin |
| Bladder | Endometrium, pre-menopausal | Liver | Prostate | Spleen |
| Bone marrow | Endometrium, post-menopausal | Lung | Rectum | Stomach |
| Breast | Epididymis | Lymph node | Seminal vesicle | Testis |
| Bronchus | Esophagus | Nasopharynx | Salivary gland | Thyroid gland |
| Caudate& cerebellum | Fallopian tube | Oral tissue | Skeletal muscle | Tonsil |
| Cervix | Gall bladder | Ovary | Small intestine | Vagina |
| Colon | Heart muscle | Pancreas | Smooth muscle | Vulva/anal |

Table S5: Basic Panel: Mapping of Original Labels to Standardized Labels across Datasets

| **Dataset Name** | **Original Label** | **Standardized Label** |
|---|---|---|
| Breast_cancer_i | OTHER | others/undefined |
| Breast_cancer_i | CD8T | CD8+ T cells |
| Breast_cancer_i | CD4T | CD4+ T cells |
| Breast_cancer_i | BCELL | B cells |
| Breast_cancer_i | TCELL | Other T cells |
| Breast_cancer_i | DC | Dendritic Cell |
| Breast_cancer_i | all others | all other cells |
| CRC | B cells | B cells |
| CRC | CD4+ T cells | CD4+ T cells |
| CRC | CD4+ T cells CD45RO+ | CD4+ T cells |
| CRC | CD4+ T cells GATA3+ | CD4+ T cells |
| CRC | CD8+ T cells | CD8+ T cells |
| CRC | CD3+ T cells | Other T cells |
| CRC | Tregs | Other T cells |
| CRC | CD11c+ DCs | Dendritic Cell |
| CRC | undefined | others/undefined |
| CRC | all others | all other cells |
| EVT | B cells | B cells |
| EVT | CD4T | CD4+ T cells |
| EVT | CD8T | CD8+ T cells |
| EVT | Treg | Other T cells |
| EVT | DC | Dendritic Cell |
| EVT | other | others/undefined |
| EVT | all others | all other cells |
| Intestine | B | B cells |
| Intestine | Plasma | B cells |
| Intestine | CD4+ T cell | CD4+ T cells |
| Intestine | CD8+ T | CD8+ T cells |
| Intestine | Treg | Other T cells |
| Intestine | DC | Dendritic Cell |
| Intestine | other | others/undefined |
| Intestine | all others | all other cells |
| Lymphoma | B | B cells |
| Lymphoma | PC | B cells |
| Lymphoma | TFH | CD4+ T cells |
| Lymphoma | CD4T | CD4+ T cells |
| Lymphoma | CD4+ T cell | CD4+ T cells |
| Lymphoma | CD4TNaive | CD4+ T cells |
| Lymphoma | TTOX | CD8+ T cells |
| Lymphoma | TTOX_exh | CD8+ T cells |

| Dataset Name | Original Label | Standardized Label |
|---|---|---|
| Lymphoma | TTOXNaive | CD8+ T cells |
| Lymphoma | Treg | Other T cells |
| Lymphoma | FDC | Dendritic Cell |
| Lymphoma | DC | Dendritic Cell |
| Lymphoma | na | others/undefined |
| Lymphoma | all others | all other cells |
| TB | B_cell | B cells |
| TB | CD4_T | CD4+ T cells |
| TB | CD8_T | CD8+ T cells |
| TB | gdT_cell | Other T cells |
| TB | Treg | Other T cells |
| TB | CD11c_DC/Mono | Dendritic Cell |
| TB | CD209_DC | Dendritic Cell |
| TB | all others | all other cells |
| cHL_MIBI | B | B cells |
| cHL_MIBI | CD4 T | CD4+ T cells |
| cHL_MIBI | CD4 Treg | CD4+ T cells |
| cHL_MIBI | CD8 T | CD8+ T cells |
| cHL_MIBI | Treg | Other T cells |
| cHL_MIBI | DC | Dendritic Cell |
| cHL_MIBI | Other | others/undefined |
| cHL_MIBI | all others | all other cells |
| cHL_CODEX | B | B cells |
| cHL_CODEX | CD4 | CD4+ T cells |
| cHL_CODEX | CD8 | CD8+ T cells |
| cHL_CODEX | Cytotoxic CD8 | CD8+ T cells |
| cHL_CODEX | TReg | Other T cells |
| cHL_CODEX | DC | Dendritic Cell |
| cHL_CODEX | Seg Artifact | others/undefined |
| cHL_CODEX | Other | others/undefined |
| cHL_CODEX | all others | all other cells |
| Bone_Marrow | B-Cells | B cells |
| Bone_Marrow | Plasma Cells | B cells |
| Bone_Marrow | Immature_B_Cell | B cells |
| Bone_Marrow | CD4+ T-Cell | CD4+ T cells |
| Bone_Marrow | CD4 Treg | CD4+ T cells |
| Bone_Marrow | CD8+ T-Cell | CD8+ T cells |
| Bone_Marrow | pDC | Dendritic Cell |
| Bone_Marrow | Artifact | others/undefined |
| Bone_Marrow | Undetermined | others/undefined |
| Bone_Marrow | Autofluorescent | others/undefined |
| Bone_Marrow | all others | all other cells |
| AML | B-Cells | B cells |
| AML | Plasma Cells | B cells |
| AML | Immature_B_Cell | B cells |
| AML | CD4+ T-Cell | CD4+ T cells |
| AML | CD4 Treg | CD4+ T cells |
| AML | CD8+ T-Cell | CD8+ T cells |
| AML | pDC | Dendritic Cell |
| AML | Artifact | others/undefined |
| AML | Undetermined | others/undefined |
| AML | Autofluorescent | others/undefined |
| AML | all others | all other cells |

Table S6: Full Panel: Mapping of Original Labels to Standardized Labels across Datasets

| Dataset Name | Original Label | Standardized Label |
|---|---|---|
| Breast_cancer_i | TUMOR_LUMINAL | others (defined but not included) |
| Breast_cancer_i | TUMOR_EMT | others (defined but not included) |
| Breast_cancer_i | TUMOR_ECADCK | others (defined but not included) |
| Breast_cancer_i | TUMOR_CK5 | others (defined but not included) |
| Breast_cancer_i | ENDO | others (defined but not included) |
| Breast_cancer_i | OTHER | artifact/not defined |
| Breast_cancer_i | NORMFIBRO | others (defined but not included) |
| Breast_cancer_i | NEUT | Granulocyte (neutrophil) |
| Breast_cancer_i | MYOFIBRO | others (defined but not included) |
| Breast_cancer_i | MYOEP | others (defined but not included) |
| Breast_cancer_i | MONODC | Dendritic cell |
| Breast_cancer_i | MONO | Macrophage |
| Breast_cancer_i | MAST | Mast cell |
| Breast_cancer_i | MACS | others (defined but not included) |
| Breast_cancer_i | FIBRO_VIMonly | others (defined but not included) |
| Breast_cancer_i | CD8T | CD8 T cell |
| Breast_cancer_i | CD4T | CD4 T cell |
| Breast_cancer_i | CAF | others (defined but not included) |
| Breast_cancer_i | BCELL | B cell |
| Breast_cancer_i | APC | others (defined but not included) |
| Breast_cancer_i | TCELL | others (defined but not included) |
| Breast_cancer_i | IMMUNEOTHER | others (defined but not included) |
| Breast_cancer_i | DC | Dendritic cell |
| Lymph_node | B | B cell |
| Lymph_node | FDC | Dendritic cell |
| Lymph_node | TTOX | CD8 T cell |
| Lymph_node | Macro | Macrophage |
| Lymph_node | na | artifact/not defined |
| Lymph_node | CD4T | CD4 T cell |
| Lymph_node | Stromal cells | others (defined but not included) |
| Lymph_node | TFH | CD4 T cell |
| Lymph_node | DC | Dendritic cell |
| Lymph_node | Granulo | Granulocyte (neutrophil) |
| Lymph_node | Treg | Treg cell |
| Lymph_node | TPR | others (defined but not included) |
| Lymph_node | NK | NK cell |
| Lymph_node | CD4TNaive | CD4 T cell |
| Lymph_node | TTOX_exh | CD8 T cell |
| Lymph_node | TTOXNaive | CD8 T cell |
| Lymph_node | MC | Macrophage |
| Lymph_node | PC | Plasma cell |
| Lymph_node | NKT | NK cell |
| cHL_MIBI | Other | artifact/not defined |
| cHL_MIBI | CD4 T | CD4 T cell |
| cHL_MIBI | Endothelial | others (defined but not included) |
| cHL_MIBI | M1 | Macrophage |
| cHL_MIBI | B | B cell |
| cHL_MIBI | M2 | M2 Macrophage |
| cHL_MIBI | NK | NK cell |
| cHL_MIBI | DC | Dendritic cell |
| cHL_MIBI | CD4 Treg | Treg cell |
| cHL_MIBI | Neutrophil | Granulocyte (neutrophil) |

| Dataset Name | Original Label | Standardized Label |
|---|---|---|
| cHL_MIBI | CD8 T | CD8 T cell |
| cHL_MIBI | Tumor | others (defined but not included) |
| cHL_CODEX | B | B cell |
| cHL_CODEX | DC | Dendritic cell |
| cHL_CODEX | NK | NK cell |
| cHL_CODEX | Monocyte | Macrophage |
| cHL_CODEX | CD4 | CD4 T cell |
| cHL_CODEX | Lymphatic | others (defined but not included) |
| cHL_CODEX | CD8 | CD8 T cell |
| cHL_CODEX | M1 | Macrophage |
| cHL_CODEX | Seg Artifact | artifact/not defined |
| cHL_CODEX | Endothelial | others (defined but not included) |
| cHL_CODEX | Mast | Mast cell |
| cHL_CODEX | Tumor | others (defined but not included) |
| cHL_CODEX | Neutrophil | Granulocyte (neutrophil) |
| cHL_CODEX | M2 | M2 Macrophage |
| cHL_CODEX | TReg | Treg cell |
| cHL_CODEX | Cytotoxic CD8 | CD8 T cell |
| cHL_CODEX | Epithelial | others (defined but not included) |
| cHL_CODEX | Other | artifact/not defined |
| CRC | granulocytes | Granulocyte (neutrophil) |
| CRC | vasculature | others (defined but not included) |
| CRC | CD4+ T cells CD45RO+ | CD4 T cell |
| CRC | tumor cells | others (defined but not included) |
| CRC | stroma | others (defined but not included) |
| CRC | CD68+CD163+ macrophages | M2 Macrophage |
| CRC | adipocytes | others (defined but not included) |
| CRC | Plasma cells | Plasma cell |
| CRC | CD8+ T cells | CD8 T cell |
| CRC | dirt | artifact/not defined |
| CRC | Tregs | Treg cell |
| CRC | CD4+ T cells | CD4 T cell |
| CRC | CD11c+ DCs | Dendritic cell |
| CRC | B cells | B cell |
| CRC | CD11b+CD68+ macrophages | Macrophage |
| CRC | smooth muscle | others (defined but not included) |
| CRC | undefined | artifact/not defined |
| CRC | tumor cells / immune cells | others (defined but not included) |
| CRC | immune cells / vasculature | others (defined but not included) |
| CRC | immune cells | others (defined but not included) |
| CRC | NK cells | NK cell |
| CRC | nerves | others (defined but not included) |
| CRC | CD68+ macrophages GzmB+ | Macrophage |
| CRC | CD68+ macrophages | Macrophage |
| CRC | lymphatics | others (defined but not included) |
| CRC | CD11b+ monocytes | Macrophage |
| CRC | CD4+ T cells GATA3+ | CD4 T cell |
| CRC | CD163+ macrophages | M2 Macrophage |
| CRC | CD3+ T cells | others (defined but not included) |
| EVT | Mac2a | M2 Macrophage |
| EVT | other | artifact/not defined |
| EVT | NK1 | NK cell |
| EVT | Fibroblasts | others (defined but not included) |
| EVT | NKT | NK cell |
| EVT | Endothelial | others (defined but not included) |

| Dataset Name | Original Label | Standardized Label |
|---|---|---|
| EVT | Myofibroblasts | others (defined but not included) |
| EVT | Mac1a | Macrophage |
| EVT | EVT1a | others (defined but not included) |
| EVT | Mac1b | Macrophage |
| EVT | CD8T | CD8 T cell |
| EVT | EVT1b | others (defined but not included) |
| EVT | Mac2c | M2 Macrophage |
| EVT | NK2 | NK cell |
| EVT | muscle | others (defined but not included) |
| EVT | NK3 | NK cell |
| EVT | EVT2 | others (defined but not included) |
| EVT | Mac2b | M2 Macrophage |
| EVT | DC | Dendritic cell |
| EVT | Glandular | others (defined but not included) |
| EVT | CD4T | CD4 T cell |
| EVT | EVT1c | others (defined but not included) |
| EVT | Placental_Mac | M2 Macrophage |
| EVT | NK4 | NK cell |
| EVT | Mast | Mast cell |
| EVT | Treg | Treg cell |

Table S7: Extended Panel: Mapping of Original Labels to Standardized Labels across Datasets

| Dataset Name | Original Label | Standardized Label |
|---|---|---|
| TB | CD163_Mac | M2 Macrophage |
| TB | CD68_Mac | Macrophage |
| TB | CD16_CD14_Mono | Macrophage |
| TB | CD4_T | CD4+ T cell |
| TB | CD14_Mono | Macrophage |
| TB | fibroblast | other (defined but not included) |
| TB | endothelial | other (defined but not included) |
| TB | Treg | Treg cell |
| TB | CD11c_DC/Mono | Dendritic cell |
| TB | imm_other | other (defined but not included) |
| TB | CD11b/c_CD206_Mac/Mono | Macrophage |
| TB | CD8_T | CD8+ T cell |
| TB | B_cell | B cell |
| TB | neutrophil | Granulocyte |
| TB | CD206_Mac | M2 Macrophage |
| TB | mast | Mast cell |
| TB | epithelial | other (defined but not included) |
| TB | giant_cell | other (defined but not included) |
| TB | gdT_cell | other (defined but not included) |
| TB | CD209_DC | Dendritic cell |
| Intestine | NK | NK cell |
| Intestine | Enterocyte | other (defined but not included) |
| Intestine | MUC1+ Enterocyte | other (defined but not included) |
| Intestine | TA | other (defined but not included) |
| Intestine | CD66+ Enterocyte | other (defined but not included) |
| Intestine | Paneth | other (defined but not included) |
| Intestine | Cycling TA | other (defined but not included) |
| Intestine | Goblet | other (defined but not included) |
| Intestine | Smooth muscle | other (defined but not included) |
| Intestine | M1 Macrophage | Macrophage |
| Intestine | Neuroendocrine | other (defined but not included) |
| Intestine | Stroma | other (defined but not included) |
| Intestine | Nerve | other (defined but not included) |
| Intestine | ICC | other (defined but not included) |
| Intestine | Lymphatic | other (defined but not included) |
| Intestine | Plasma | Plasma cell |
| Intestine | CD4+ T cell | CD4+ T cell |
| Intestine | M2 Macrophage | M2 Macrophage |
| Intestine | DC | Dendritic cell |
| Intestine | B | B cell |
| Intestine | CD57+ Enterocyte | other (defined but not included) |
| Intestine | CD8+ T | CD8+ T cell |
| Intestine | Neutrophil | Granulocyte |
| Intestine | Endothelial | other (defined but not included) |
| Intestine | Cycling TA | other (defined but not included) |
| Intestine | CD7+ Immune | other (defined but not included) |
| Bone Marrow | SEC | other (defined but not included) |
| Bone Marrow | Intermediate Myeloid | other (defined but not included) |
| Bone Marrow | Erythroblast | other (defined but not included) |
| Bone Marrow | Mature Myeloid | other (defined but not included) |
| Bone Marrow | Erythroid | other (defined but not included) |
| Bone Marrow | Plasma Cells | Plasma cell |

| Dataset Name | Original Label | Standardized Label |
|---|---|---|
| Bone Marrow | NPM1 Mutant Blast | other (defined but not included) |
| Bone Marrow | CD8+ T-Cell | CD8+ T cell |
| Bone Marrow | B-Cells | B cell |
| Bone Marrow | Early Myeloid Progenitor | other (defined but not included) |
| Bone Marrow | Adipocyte | other (defined but not included) |
| Bone Marrow | CD4+ T-Cell | CD4+ T cell |
| Bone Marrow | Macrophages | Macrophage |
| Bone Marrow | AEC | other (defined but not included) |
| Bone Marrow | Immature_B_Cell | B cell |
| Bone Marrow | GATA1pos_Mks | other (defined but not included) |
| Bone Marrow | Monocytes | Macrophage |
| Bone Marrow | pDC | Dendritic cell |
| Bone Marrow | Endosteal | other (defined but not included) |
| Bone Marrow | Adipo-MSC | other (defined but not included) |
| Bone Marrow | Non-Classical Monocyte | Macrophage |
| Bone Marrow | GMP | other (defined but not included) |
| Bone Marrow | HSPC | other (defined but not included) |
| Bone Marrow | THY1+ MSC | other (defined but not included) |
| Bone Marrow | VSMC | other (defined but not included) |
| Bone Marrow | CD34+ CD61+ | other (defined but not included) |
| Bone Marrow | GATA1neg_Mks | other (defined but not included) |
| Bone Marrow | SPINK2+ HSPC | other (defined but not included) |
| Bone Marrow | Artifact | artifact/not defined |
| Bone Marrow | Undetermined | artifact/not defined |
| Bone Marrow | Autofluorescent | artifact/not defined |
| Bone Marrow | CD44+ Undetermined | other (defined but not included) |
| Bone Marrow | MEP/Early Erythroblast | other (defined but not included) |
| Bone Marrow | GMP/Myeloblast | other (defined but not included) |
| Bone Marrow | HSC | other (defined but not included) |
| Bone Marrow | CLP | other (defined but not included) |
| Bone Marrow | Schwann Cells | other (defined but not included) |

Table S8: Structure Panel: Mapping of Original Labels to Standardized Labels across Datasets

| Dataset Name | Original Label | Standardized Label |
|---|---|---|
| CRC | granulocytes | all the other cell types |
| CRC | vasculature | endothelial |
| CRC | CD4+ T cells CD45RO+ | all the other cell types |
| CRC | tumor cells | tumor |
| CRC | stroma | stroma |
| CRC | CD68+CD163+ macrophages | all the other cell types |
| CRC | adipocytes | all the other cell types |
| CRC | plasma cells | all the other cell types |
| CRC | CD8+ T cells | all the other cell types |
| CRC | dirt | undefined |
| CRC | Tregs | all the other cell types |
| CRC | CD4+ T cells | all the other cell types |
| CRC | CD11c+ DCs | all the other cell types |
| CRC | B cells | all the other cell types |
| CRC | CD11b+CD68+ macrophages | all the other cell types |
| CRC | smooth muscle | smooth muscle |
| CRC | undefined | undefined |
| CRC | tumor cells / immune cells | tumor |
| CRC | immune cells / vasculature | endothelial |
| CRC | immune cells | all the other cell types |
| CRC | NK cells | all the other cell types |
| CRC | nerves | all the other cell types |
| CRC | CD68+ macrophages GzmB+ | all the other cell types |
| CRC | CD68+ macrophages | all the other cell types |
| CRC | lymphatics | smooth muscle |
| CRC | CD11b+ monocytes | all the other cell types |
| CRC | CD4+ T cells GATA3+ | all the other cell types |
| CRC | CD163+ macrophages | all the other cell types |
| CRC | CD3+ T cells | all the other cell types |
| cHL_CODEX | B | all the other cell types |
| cHL_CODEX | DC | all the other cell types |
| cHL_CODEX | NK | all the other cell types |
| cHL_CODEX | Monocyte | all the other cell types |
| cHL_CODEX | CD4 | all the other cell types |
| cHL_CODEX | Lymphatic | smooth muscle |
| cHL_CODEX | CD8 | all the other cell types |
| cHL_CODEX | M1 | all the other cell types |
| cHL_CODEX | Seg Artifact | undefined |
| cHL_CODEX | Endothelial | endothelial |
| cHL_CODEX | Mast | all the other cell types |
| cHL_CODEX | Tumor | tumor |
| cHL_CODEX | Neutrophil | all the other cell types |
| cHL_CODEX | M2 | all the other cell types |
| cHL_CODEX | TReg | all the other cell types |
| cHL_CODEX | Cytotoxic CD8 | all the other cell types |
| cHL_CODEX | Epithelial | epithelial |
| cHL_CODEX | Other | undefined |
| Breast Cancer | TUMOR_LUMINAL | tumor |
| Breast Cancer | TUMOR_EMT | tumor |
| Breast Cancer | TUMOR_ECADCK | tumor |
| Breast Cancer | TUMOR_CK5 | tumor |
| Breast Cancer | ENDO | endothelial |

| Dataset Name | Original Label | Standardized Label |
|---|---|---|
| Breast Cancer | OTHER | undefined |
| Breast Cancer | NORMFIBRO | stroma |
| Breast Cancer | NEUT | all the other cell types |
| Breast Cancer | MYOFIBRO | smooth muscle |
| Breast Cancer | MYOEP | epithelial |
| Breast Cancer | MONODC | all the other cell types |
| Breast Cancer | MONO | all the other cell types |
| Breast Cancer | MAST | all the other cell types |
| Breast Cancer | MACS | undefined |
| Breast Cancer | FIBRO_VIMonly | stroma |
| Breast Cancer | CD8T | all the other cell types |
| Breast Cancer | CD4T | all the other cell types |
| Breast Cancer | CAF | stroma |
| Breast Cancer | BCELL | all the other cell types |
| Breast Cancer | APC | all the other cell types |
| Breast Cancer | TCELL | all the other cell types |
| Breast Cancer | IMMUNEOTHER | all the other cell types |
| Breast Cancer | DC | all the other cell types |
| EVT | Mac2a | all the other cell types |
| EVT | other | undefined |
| EVT | NK1 | all the other cell types |
| EVT | Fibroblasts | stroma |
| EVT | NKT | all the other cell types |
| EVT | Endothelial | endothelial |
| EVT | Myofibroblasts | smooth muscle |
| EVT | Mac1a | all the other cell types |
| EVT | EVT1a | all the other cell types |
| EVT | Mac1b | all the other cell types |
| EVT | CD8T | all the other cell types |
| EVT | EVT1b | all the other cell types |
| EVT | Mac2c | all the other cell types |
| EVT | NK2 | all the other cell types |
| EVT | muscle | smooth muscle |
| EVT | NK3 | all the other cell types |
| EVT | EVT2 | all the other cell types |
| EVT | Mac2b | all the other cell types |
| EVT | DC | all the other cell types |
| EVT | Glandular | epithelial |
| EVT | CD4T | all the other cell types |
| EVT | EVT1c | all the other cell types |
| EVT | Placental_Mac | all the other cell types |
| EVT | NK4 | all the other cell types |
| EVT | Mast | all the other cell types |
| EVT | Treg | all the other cell types |

Table S9: Nerve Panel: Mapping of Original Labels to Standardized Labels across Datasets

| **Dataset Name** | **Original Label** | **Standardized Label** |
|---|---|---|
| Tonsil_Unlabeled | Innate | all the other cell types |
| Tonsil_Unlabeled | PDPN | all the other cell types |
| Tonsil_Unlabeled | Endothelial | all the other cell types |
| Tonsil_Unlabeled | B cell | all the other cell types |
| Tonsil_Unlabeled | T cell | all the other cell types |
| Tonsil_Unlabeled | Squamous epithelial | all the other cell types |
| Tonsil_Unlabeled | Stromal | all the other cell types |
| Tonsil_Unlabeled | smooth muscle | all the other cell types |
| Tonsil_Unlabeled | Plasma Cell | all the other cell types |
| Tonsil_Unlabeled | Nerve | nerve |
| CRC | granulocytes | all the other cell types |
| CRC | vasculature | all the other cell types |
| CRC | CD4+ T cells CD45RO+ | all the other cell types |
| CRC | tumor cells | all the other cell types |
| CRC | stroma | all the other cell types |
| CRC | CD68+CD163+ macrophages | all the other cell types |
| CRC | adipocytes | all the other cell types |
| CRC | plasma cells | all the other cell types |
| CRC | CD8+ T cells | all the other cell types |
| CRC | dirt | undefined |
| CRC | Tregs | all the other cell types |
| CRC | CD4+ T cells | all the other cell types |
| CRC | CD11c+ DCs | all the other cell types |
| CRC | B cells | all the other cell types |
| CRC | CD11b+CD68+ macrophages | all the other cell types |
| CRC | smooth muscle | all the other cell types |
| CRC | undefined | undefined |
| CRC | tumor cells / immune cells | all the other cell types |
| CRC | immune cells / vasculature | all the other cell types |
| CRC | immune cells | all the other cell types |
| CRC | NK cells | all the other cell types |
| CRC | nerves | nerve |
| CRC | CD68+ macrophages GzmB+ | all the other cell types |
| CRC | CD68+ macrophages | all the other cell types |
| CRC | lymphatics | all the other cell types |
| CRC | CD11b+ monocytes | all the other cell types |
| CRC | CD4+ T cells GATA3+ | all the other cell types |
| CRC | CD163+ macrophages | all the other cell types |
| CRC | CD3+ T cells | all the other cell types |
| Intestine | NK | all the other cell types |
| Intestine | Enterocyte | all the other cell types |
| Intestine | MUC1+ Enterocyte | all the other cell types |
| Intestine | TA | all the other cell types |
| Intestine | CD66+ Enterocyte | all the other cell types |
| Intestine | Paneth | all the other cell types |
| Intestine | Cycling TA | all the other cell types |
| Intestine | Goblet | all the other cell types |
| Intestine | Smooth muscle | all the other cell types |
| Intestine | M1 Macrophage | all the other cell types |
| Intestine | Neuroendocrine | nerve |
| Intestine | Stroma | all the other cell types |
| Intestine | Nerve | nerve |

| **Dataset Name** | **Original Label** | **Standardized Label** |
|---|---|---|
| Intestine | ICC | all the other cell types |
| Intestine | Lymphatic | all the other cell types |
| Intestine | Plasma | all the other cell types |
| Intestine | CD4+ T cell | all the other cell types |
| Intestine | M2 Macrophage | all the other cell types |
| Intestine | DC | all the other cell types |
| Intestine | B | all the other cell types |
| Intestine | CD57+ Enterocyte | all the other cell types |
| Intestine | CD8+ T | all the other cell types |
| Intestine | M2 Macrophage | all the other cell types |
| Intestine | Neutrophil | all the other cell types |
| Intestine | Endothelial | all the other cell types |
| Intestine | Cycling TA | all the other cell types |
| Intestine | CD7+ Immune | all the other cell types |
| Intestine | Undefined (NaN) | undefined |

#### 2 Supplementary figures

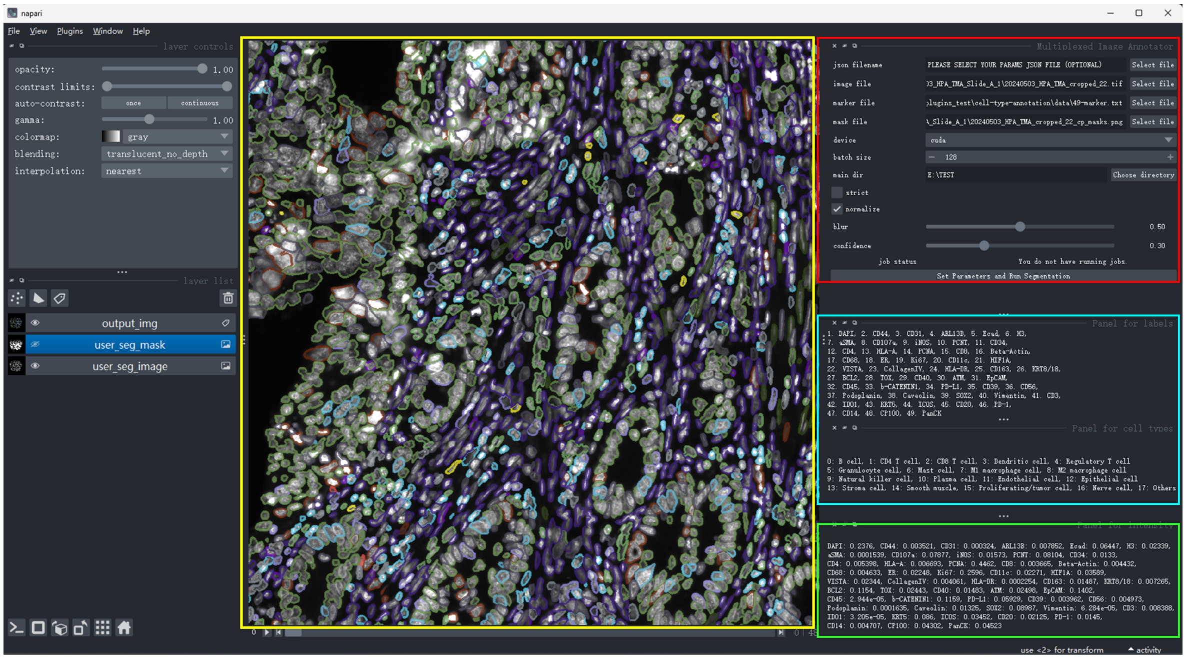


Figure S1: Napari-based visualization of our cell type annotator. The main Napari build-in viewer (yellow box) shows the original tissue image, cell segmentation, and our annotation results in three layers; the panel in red box is for user to required file paths and hyperparameters; below, the panel in cyan lists marker and cell type names; when the user clicks any cells on the cell segmentation layer in the main viewer, the green panel will show the cell-level marker expression intensity of that cell.

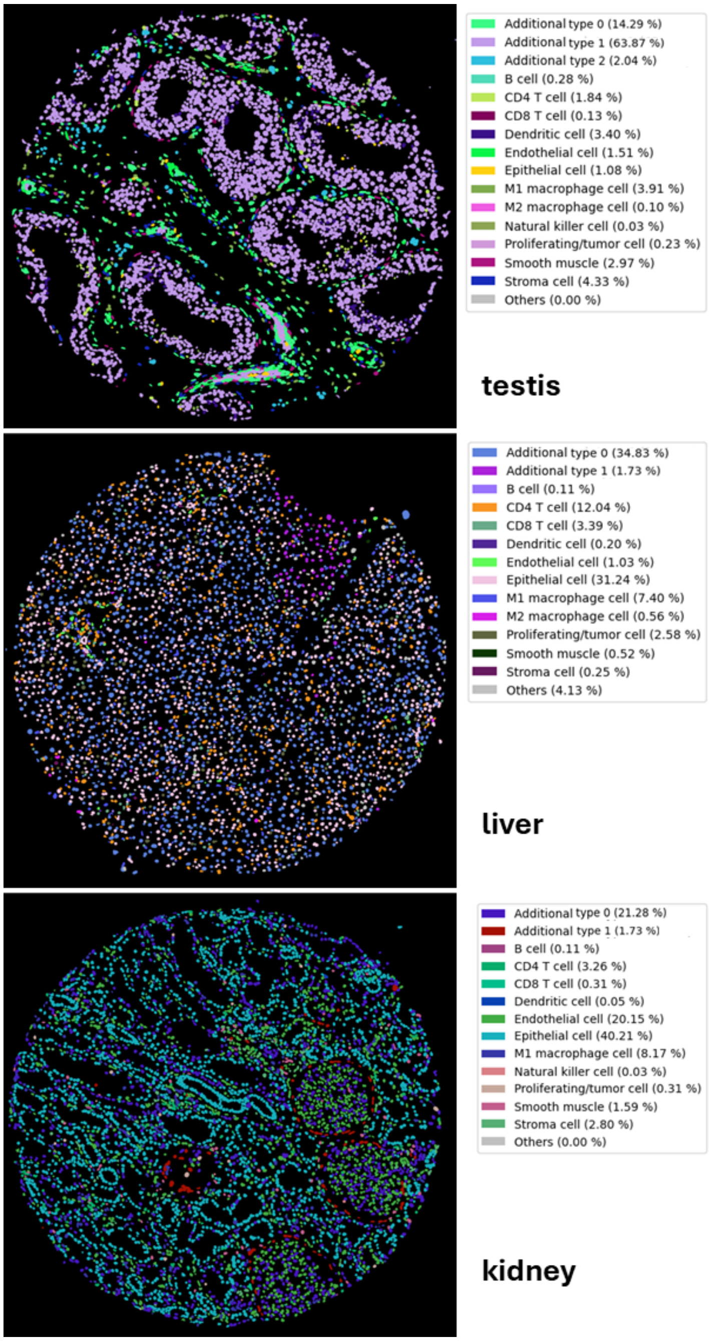


Figure S2: RIBCA enables the identification of tissue-specific cell types through unsupervised clustering of cells that were not previously annotated to 18 known cell types. We identified interstitial cells (additional subtypes 0 and 1) and epididymal duct (additional subtype 2) in testis tissue; kupffer cell (additional subtype 1) in liver tissue.

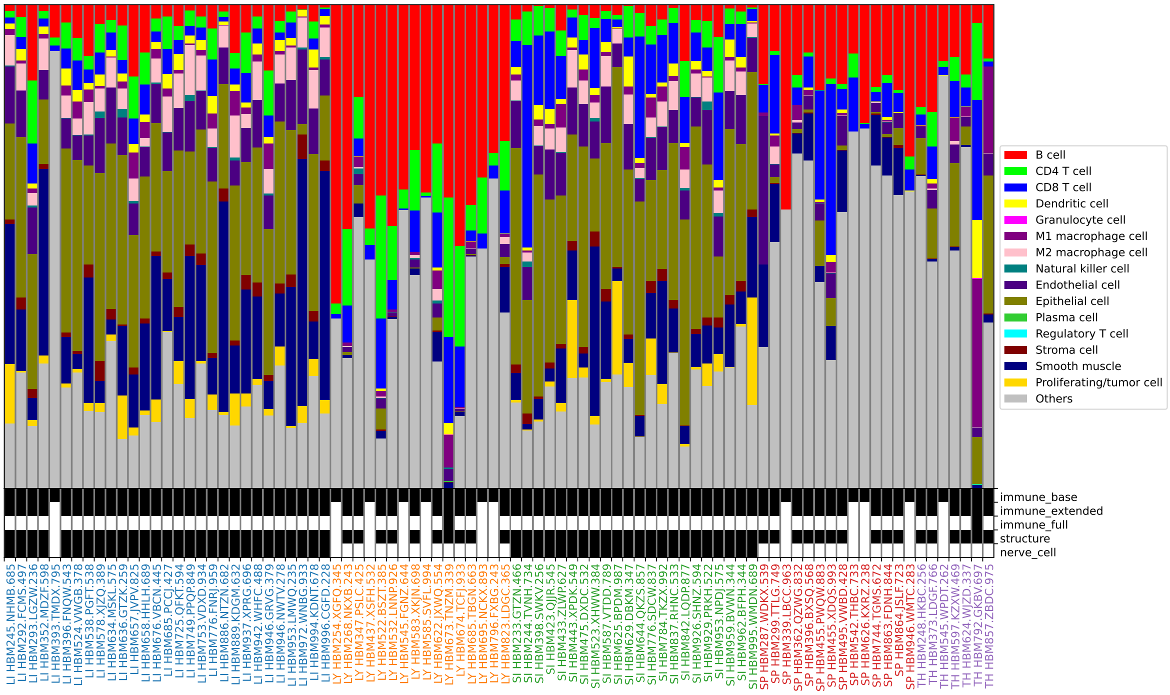


Figure S3: Analysis of individual HuBMAP CODEX datasets. The tissue and the HuBMAP dataset ID are shown along the bottom (the corresponding datasets can be viewed and downloaded from the HuBMAP portal using the IDs). Rows labeled with the names of RIBCA panels show in black those datasets that have sufficient markers to run that panel (note that not all markers are needed since some can be imputed). The fraction of cell types assigned are shown. Note the relative consistency of results within a tissue for those datasets with at least the immune_extended and structure panels (and the fluctuations among the others). The averages for those are shown in Figure 3G.

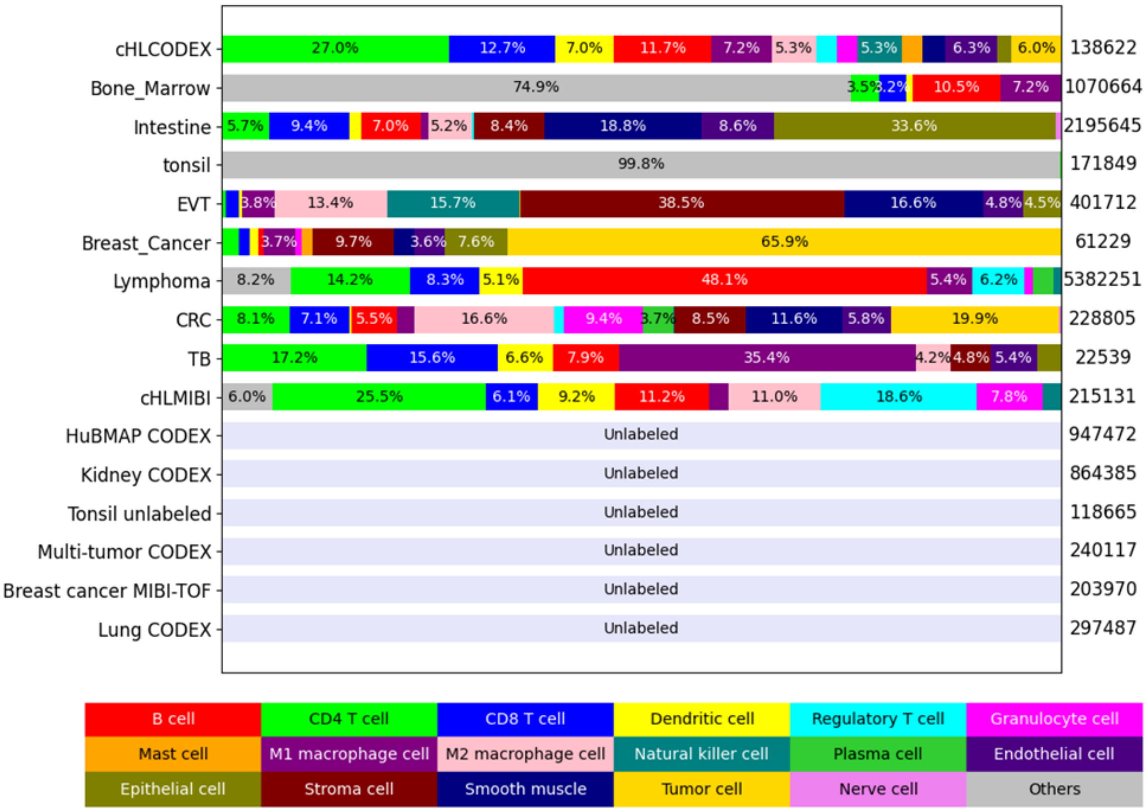


Figure S4: Statistics (cell type distributions and number of cells) of multiple public multiplexed imaging datasets used to curate our dataset to train the models. We reassigned existing annotations to 18 consistent cell types. This set of cell types is at an appropriate granularity and compatible with most existing annotations.

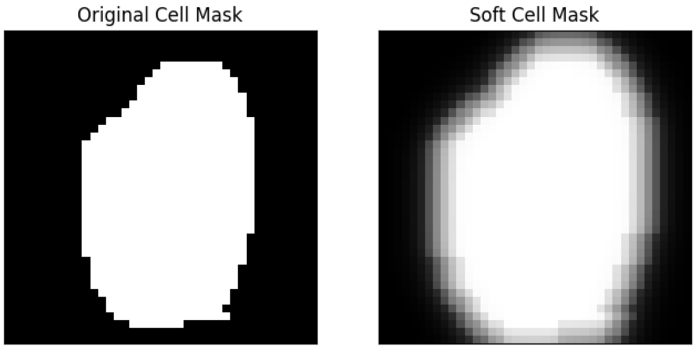


Figure S5: An example of applying soft-masking to a cell segmentation mask.

### References


[1] S. Bandyopadhyay, M. P. Duffy, K. J. Ahn, J. H. Sussman, M. Pang, D. Smith, G. Duncan, I. Zhang, J. Huang, Y. Lin, et al. Mapping the cellular biogeography of human bone marrow niches using single-cell transcriptomics and proteomic imaging. *Cell*, 187(12):3120–3140, 2024.

[2] S. Black, D. Phillips, J. W. Hickey, J. Kennedy-Darling, V. G. Venkataraaman, N. Samusik, Y. Goltsev, C. M. Schürch, and G. P. Nolan. Codex multiplexed tissue imaging with dna-conjugated antibodies. *Nature protocols*, 16(8):3802–3835, 2021.

[3] S. Greenbaum, I. Averbukh, E. Soon, G. Rizzuto, A. Baranski, N. F. Greenwald, A. Kagel, M. Bosse, E. G. Jaswa, Z. Khair, et al. A spatially resolved timeline of the human maternal–fetal interface. *Nature*, 619(7970):595–605, 2023.

[4] J. Hansen, R. Sealfon, R. Menon, M. T. Eadon, B. B. Lake, B. Steck, K. Anjani, S. Parikh, T. K. Sigdel, G. Zhang, et al. A reference tissue atlas for the human kidney. *Science advances*, 8(23): eabn4965, 2022.

[5] J. W. Hickey, W. R. Becker, S. A. Nevins, A. Horning, A. E. Perez, C. Zhu, B. Zhu, B. Wei, R. Chiu, D. C. Chen, et al. Organization of the human intestine at single-cell resolution. *Nature*, 619(7970):572–584, 2023.

[6] L. Keren, M. Bosse, D. Marquez, R. Angoshtari, S. Jain, S. Varma, S.-R. Yang, A. Kurian, D. Van Valen, R. West, et al. A structured tumor-immune microenvironment in triple negative breast cancer revealed by multiplexed ion beam imaging. *Cell*, 174(6):1373–1387, 2018.

[7] E. F. McCaffrey, M. Donato, L. Keren, Z. Chen, A. Delmastro, M. B. Fitzpatrick, S. Gupta, N. F. Greenwald, A. Baranski, W. Graf, et al. The immunoregulatory landscape of human tuberculosis granulomas. *Nature immunology*, 23(2):318–329, 2022.

[8] T. Risom, D. R. Glass, I. Averbukh, C. C. Liu, A. Baranski, A. Kagel, E. F. McCaffrey, N. F. Greenwald, B. Rivero-Gutiérrez, S. H. Strand, et al. Transition to invasive breast cancer is associated with progressive changes in the structure and composition of tumor stroma. *Cell*, 185 (2):299–310, 2022.

[9] T. Roider, M. A. Baertsch, D. Fitzgerald, H. Voehringer, B. J. Brinkmann, F. Czernilofsky, M. Knoll, L. Llaó-Cid, A. Mathioudaki, B. Faßbender, et al. Multimodal and spatially resolved profiling identifies distinct patterns of t cell infiltration in nodal b cell lymphoma entities. *Nature Cell Biology*, pages 1–12, 2024.

[10] C. M. Schürch, S. S. Bhate, G. L. Barlow, D. J. Phillips, L. Noti, I. Zlobec, P. Chu, S. Black, J. Demeter, D. R. McIlwain, et al. Coordinated cellular neighborhoods orchestrate antitumoral immunity at the colorectal cancer invasive front. *Cell*, 182(5):1341–1359, 2020.

[11] M. Shaban, Y. Bai, H. Qiu, S. Mao, J. Yeung, Y. Y. Yeo, V. Shanmugam, H. Chen, B. Zhu, J. L. Weirather, et al. Maps: Pathologist-level cell type annotation from tissue images through machine learning. *Nature Communications*, 15(1):28, 2024.
